# Computational Structure Modeling, Functional Characterization, and Identification of Potential Inhibitors for the cyclic-di-GMP Mediated Biofilm Forming Membrane Protein in *Vibrio cholerae*

**DOI:** 10.64898/2026.05.26.728059

**Authors:** Md. Nahid Hasan Joy, Md Kazy Ebnul Hasan, Md. Sanowar Hossan, Md. Mahedi Hasan Sourov, Shahad Shahriar, Md. Faruk Hasan, Amit Kumar Dutta, Md. Enamul Haque

**Affiliations:** Department of Biotechnology and Genetic Engineering, Gopalganj Science and Technology University, Gopalganj, 8105, Bangladesh; Department of Genetic Engineering and Biotechnology, University of Rajshahi, Rajshahi, 6205, Bangladesh; Department of Zoology, Faculty of Biological Sciences, University of Rajshahi, Rajshahi 6205, Bangladesh; Department of Microbiology, University of Rajshahi, Rajshahi, 6205, Bangladesh; Department of Computer Science and Mathematics, Bangladesh Agricultural University, Mymensingh, 2200, Bangladesh

**Author notes:** Corresponding author: Dr. Md. Enamul Haque Assistant Professor, Department of Biotechnology and Genetic Engineering Gopalganj Science and Technology University Gopalganj-8105, Bangladesh.

**Keywords:** Biofilm formation, c-di-GMP, Diguanylate cyclase, hypothetical protein TYC33605.1, *Vibrio cholerae*

## Abstract

Cholera, caused by *Vibrio cholerae*, continues to pose a serious global public health challenge, with its impact worsened by rising antibiotic resistance associated with bacterial biofilm formation. This study reveals the role of the hypothetical protein (HP) TYC33605.1 in cyclic-di-GMP (c-di-GMP)-mediated biofilm regulation and identifies natural inhibitors that disrupt this mechanism. Functional annotation revealed TYC33605.1 as a membrane-associated diguanylate cyclase (DGC) with GGDEF and sensory domains, critical for c-di-GMP synthesis and biofilm persistence. Homology modelling and molecular dynamics (MD) simulations validated its stable 3D structure (C-score: −1.22, Ramachandran favoured regions: 91.1%) and dynamic behaviour (average RMSD: 8.55 Å). Virtual molecular docking screening of 1,092 natural compounds identified Luteolin (CID 5280445) and Sativanone (CID 13886678) as top candidates, exhibiting strong binding affinities (−9.1 and −9.0 kcal/mol, respectively) and forming hydrogen bonds, π-cation, and hydrophobic interactions with key residues (Glu293, Arg364, Ala176). MD simulations (100 ns) confirmed complex stability, with Luteolin and Sativanone showing lower RMSD fluctuations (7.78 Å and 8.13 Å) compared to the control and apoprotein. The ADME/Tox profiles highlighted favourable pharmacodynamics (PD), pharmacokinetics (PK), high gastrointestinal absorption, no hepatotoxicity, and drug-likeness (Lipinski compliance). Principal component, probability density function, and binding free energy analyses underscore ligand-induced conformational stability. This study proposes the molecular characterisation of the HP and the bioactive compounds Luteolin and Sativanone as promising inhibitors targeting TYC33605.1, offering a novel strategy to combat biofilm-mediated antibiotic resistance and a framework for analogous antimicrobial discovery in *Vibrio cholerae*.

## 1. Introduction

Cholera, an acute diarrheal disease caused by the gram-negative pathogenic bacterium *Vibrio cholerae* (Montero et al. 2023), remains a major global health concern. According to the cholera outbreak report 2024 by the World Health Organization, from January to 24 November, a total of 733,956 cholera cases and 5,162 cholera-related deaths were reported in 33 countries (five WHO regions). Compared to the same period in 2023, the statistics for November 2024 showed a 37% increase in cases and a 27% rise in the death rate (WHO report. 2024). Among the numerous strains of *Vibrio cholerae*, serogroups O1 and O139 are particularly responsible for epidemic and pandemic cholera, whereas other serogroups, collectively termed non-O1/non-O139, typically cause milder or asymptomatic infections (Taylor et al. 1987; Echazarreta and Klose 2019). Human infection begins with the consumption of infected foods or drinks, allowing *Vibrio cholerae* to colonize in the small intestine using toxin-co-regulated pilus (TCP) and other adhesion processes (Almagro-Moreno et al. 2015). Once inside the host, it secretes cholera toxin (CT), which subsequently disrupts ion balance, leading to profuse watery diarrhea (“rice water stool”). Consequently, the resulting dehydration can cause hypovolemic shock, with a mortality rate that can reach up to 50% in untreated cases (Thiagarajah and Verkman 2003; Matz and Kjelleberg 2005).

*Vibrio cholerae* depends on biofilm formation for environmental persistence and intestinal colonization. Biofilms are organized assemblies of microorganisms embedded within a self-produced matrix of extracellular polymeric substances (EPS) (Liu et al. 2022), which provides protection against environmental stresses and significantly elevates resistance to existing antibiotics, such as doxycycline, ciprofloxacin, and erythromycin (Silva and Benitez 2016; Seeber et al. 2019). The formation of biofilms and the regulation of virulence are governed by two key mechanisms: quorum sensing (QS) systems and cyclic di-GMP (c-di-GMP) signaling pathways, and these systems facilitate cell-to-cell communication and coordinate gene expression in response to changes in cellular density, particularly under low-density conditions (Chen and Michel 1998; Lim et al. 2006; Srivastava et al. 2013; Teschler et al. 2015). In *Vibrio cholerae*, the QS system is initiated by two types of autoinducers, namely cholera autoinducer-1 (CAI-1) and autoinducer-2 (AI-2), which generate chemical signals that are detected by their respective inner membrane receptors, CqsS and LuxP/Q (Rutherford et al. 2011). These receptors function as both kinases and phosphatases, enabling them to activate or deactivate downstream signaling pathways and relay information to the transcriptional regulator LuxO (Zhu et al. 2002; Papenfort and Bassler 2016). The formation of biofilms is further influenced by secondary messengers like cyclic di-GMP (c-di-GMP). In *Vibrio cholerae*, the transition between motile and sessile lifestyles is regulated by the level of increased c-di-GMP (Chen and Michel 1998; Lim et al. 2006; Srivastava et al. 2013). The initial surface attachment, cellular proliferation, and biofilm matrix development are regulated by c-di-GMP (Pederson et al. 2018). The synthesis of c-di-GMP is initiated by GGDEF domain-containing DGCs, and its degradation is carried out by EAL or HD-GYP domain-containing phosphodiesterases (PDEs) (Ryjenkov et al. 2005). Owing to their critical roles in bacterial physiology, QS systems and DGC-associated second messengers have been recognized as promising targets for novel antimicrobial drug development, particularly in response to the growing challenge of antibiotic resistance. Unlike conventional antibiotics that target essential genes, interference with QS signaling pathways circumvents the imposition of selective pressure on bacterial populations, thereby reducing the emergence of resistant strains. Over the past two decades, numerous natural quorum-sensing inhibitors (QSIs) have been discovered across a range of organisms, alongside the development of various synthetic QSIs in research laboratories (Kalia 2013; Rampioni et al. 2014). Compounds such as quercetin, naringenin, and tryptanthrin have been shown to target LuxO, a central regulator of QS, resulting in the suppression of biofilm-related gene expression and inhibition of QS-regulated phenotypes (Narendrakumar et al. 2019; Saha et al. 2023).

In contrast, the discovery of inhibitors targeting cyclic dinucleotide-based signaling pathways has advanced at a relatively slow pace. To date, only a limited number of bioactive compounds have been identified and characterized as inhibitors of cyclic dinucleotide signaling (Opoku-Temeng et al. 2016). Cranberry extract and pentacyclic triterpenoids have been shown to modulate intracellular c-di-GMP levels, thereby leading to the inhibition of biofilm formation (Pederson et al. 2018; Bhattacharya et al. 2020). Since phytochemicals are secondary metabolites produced by plants, they serve protective roles and possess therapeutic properties against biofilm formation (Suresh and Abraham 2020). Bioactive compounds are favored over synthetic ones in drug discovery due to their lower toxicity, structural diversity, chemical novelty, abundance, and bioactivity. Additionally, they offer promising therapeutic properties, particularly in addressing antibiotic resistance to developing an effective drug (Ahmed et al. 2024). This study aims to characterize an HP (accession ID: TYC33605.1) of *Vibrio cholerae* and identify promising bioactive compounds that may interact with the HP and lead to the inhibition of biofilm formation. This study utilized comprehensive bioinformatics tools regarding characterization of the HP and identified that it belongs to the DGC family, which might be involved in promoting biofilm formation in *Vibrio cholerae* via the secondary messenger c-di-GMP.

The findings of this research offer valuable insights into an underexplored regulatory protein involved in cholera pathogenesis. This research illuminates the molecular mechanisms of c-di-GMP signaling-mediated biofilm activity through the functional annotation of the HP TYC33605.1, simultaneously paving the way for the development of eco-friendly and safe therapeutics to combat *Vibrio cholerae*-associated antibiotic resistance, a public health concern.

## 2. Materials and methods

### 2.1. Sequence Acquisition and Similarity Assessment

The HP sequence was retrieved from the National Center for Biotechnology Information (NCBI) database with the accession ID TYC33605.1 and formatted in FASTA. The BLASTp tool (Johnson et al. 2008) was utilized to perform a homology search against known protein databases of the NCBI protein database (https://www.ncbi.nlm.nih.gov/) and the SwissProt database (Boeckmann et al. 2003) to discover homologous proteins with structural and putative functional similarities. This initial step enabled a preliminary prediction of the potential function of the uncharacterized protein. For functional annotation and structural prediction, a 461-amino-acid FASTA sequence was analyzed using various software tools listed in **Supplement Table 1**. Notably, no three-dimensional structure for this protein was found in the Protein Data Bank (PDB). Therefore, this study also involved predicting and validating the 3D structure of the protein through computational modeling. The full workflow of the investigation is illustrated in **Fig 1**.

**Fig 1:**
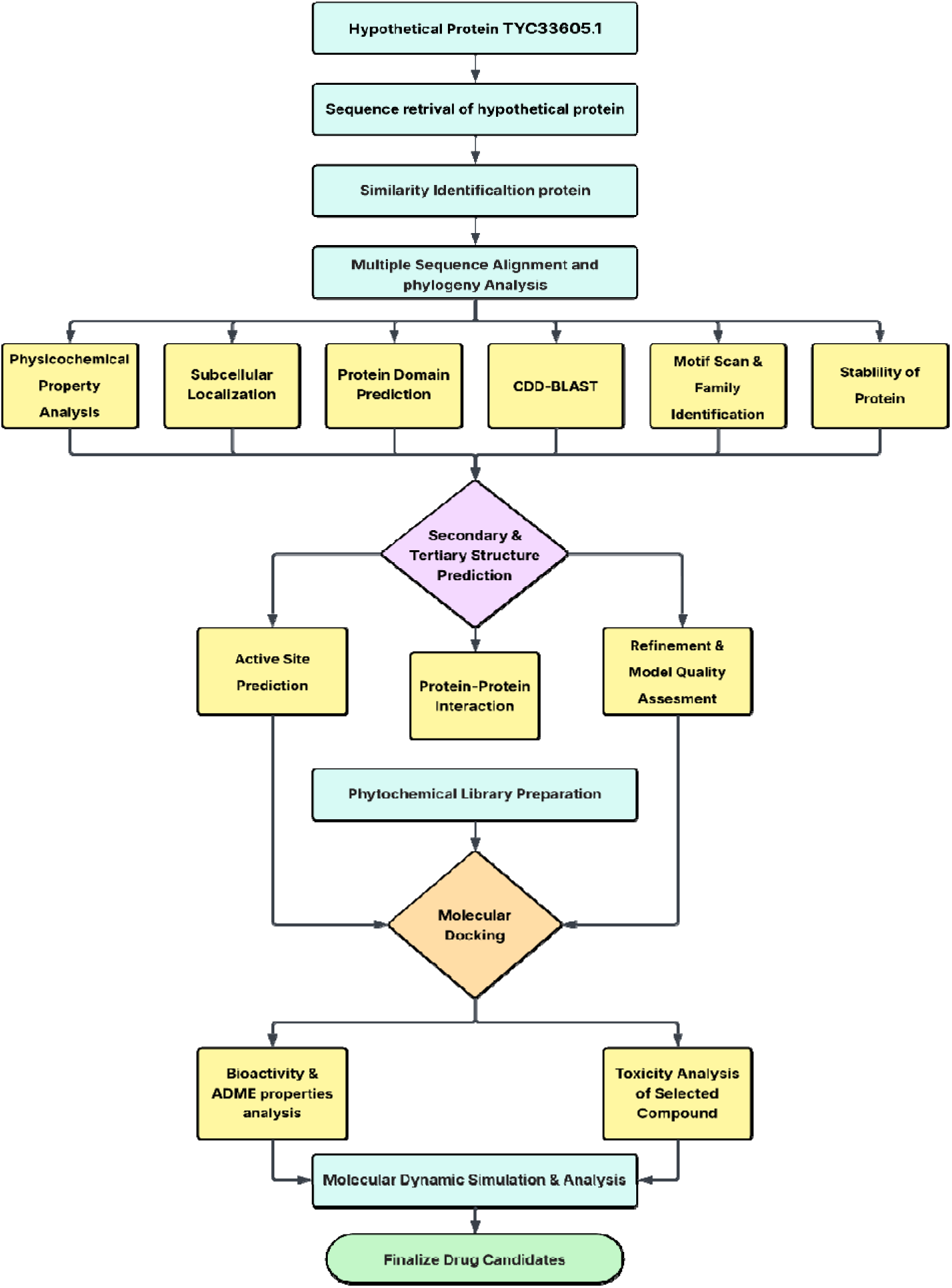
Workflow of the study outlining sequence retrieval, functional annotation, homology modeling, inhibitor screening, and validation with MD simulation analysis.

### 2.2. Multiple Sequence Alignment and Phylogenetic Analysis

Multiple sequence alignment has been done using Clustal Omega (https://www.ebi.ac.uk/Tools/msa/clustalo/) (Baxevanis et al. 2010), a program designed for rapid and precise alignment of amino acid or nucleotide sequences. The query protein’s FASTA sequence (TYC33605.1) and homologous protein sequences (AAF96459.1, WP_171356651.1, NOE85880.1, HAS7850826.1, WP_000350041.1, MVB15607.1, EKM9578803.1, WP_331305827.1, and WP_053032761.1) were used for alignment.Multiple sequence alignment has significance for predicting protein structure and function, explaining evolutionary relationships, and conducting comparative analyses. Accuracy, scalability for many proteins, and the ability to evaluate proteins with diverse domain patterns have been significantly improved by recent advancements in alignment algorithms (Katoh and Standley 2013). We used MEGA X (Molecular Evolutionary Genetics Analysis) software (Kumar et al. 2018) to construct a phylogenetic tree from the aligned sequences for phylogenetic analysis. The resultant tree file was visualized and annotated using the Interactive Tree of Life (iTOL) web server (https://itol.embl.de/) (Letunic and Bork 2021), which provides an accessible platform for the editing, analysis, and display of phylogenetic trees.

### 2.3. Physicochemical Property and Subcellular Localization Analysis

The physicochemical properties of the HP were assessed with the use of the ProtParam server, which may be found on the ExPASy website at https://web.expasy.org/protparam/ (Gasteiger et al. 2003). To understand the structural and molecular functional properties of the protein, evaluations were taken of its molecular weight (Mw), amino acid and atomic composition, number of basic (Arg + Lys) and acidic (Asp + Glu) residues, estimated half-life, instability index, theoretical isoelectric point (pI), aliphatic index, extinction coefficient, and grand average of hydropathicity (GRAVY). The subcellular location of a protein frequently reflects its biological role regarding binding with specific proteins; thus, predicting localization from its sequence can yield significant insights into its potential function (Yachdav et al. 2014). The CELLO server (Yu et al. 2014) was utilized to predict the subcellular localization of the target HP. Results were cross-validated for accuracy using additional tools such as PSORTb (Yu et al. 2010), PSLpred (Bhasin et al. 2005), and SOSUIGramN (Imai et al. 2008). Additionally, transmembrane topology was assessed utilizing the TMHMM 2.0 (Möller et al. 2001), HMMTOP (Tusnády and Simon 2001), and CCTOP (Dobson et al. 2015) web programs.

### 2.4. CDD-BLAST and Protein Domain Motif Prediction

Pfam is a reputable database recognized for its ability to accurately identify protein domains and families, offering extensive coverage (Sonnhammer et al. 1997; Finn et al. 2006; Mistry et al. 2021). The entries are categorized into two primary types: Pfam-A and Pfam-B. Entries in Pfam-A undergo meticulous curation, featuring a seed alignment that consists of a representative subset of sequences, complemented by a comprehensive alignment produced via profile Hidden Markov Model (HMM) searches against primary sequence databases. On the other hand, records from Pfam-B are made directly from the ProDom database and are shown by a single alignment, without any editing by a user (Bru et al. 2005).

Protein domains are distinct molecular evolution units that are often associated with specific molecular and cellular functions of the query protein sequence. Performing an inquiry in NCBI’s Conserved Domain Database (CDD) (https://www.ncbi.nlm.nih.gov/Structure/cdd/wrpsb.cgi) for the study of conserved domains. To effectively evaluate a set of pre-computed position-specific scoring matrices utilizing a protein search, the method employs reversed position-specific BLAST (RPSBLAST) alongside a variant of position-specific iterative BLAST (PSI-BLAST) (Marchler-Bauer et al. 2015). Performing a motif scan means identifying every known motif in a given sequence. It is common practice to use protein sequence motifs for protein function prediction and as family-specific protein IDs. The Genome Net server (https://www.genome.jp/tools/motif/) was used to search for protein motifs using Motif (Kanehisa et al. 2002).

### 2.5. Family, superfamily, fold, and coil identification

The InterProScan server (Hunter et al. 2009) was employed for the analysis and classification of protein families and superfamilies. The PFP-FunD SeqE server (Shen and Chou 2009) was used for recognizing protein folding patterns. The COILS server (Lupas et al. 1991) was employed for detecting coiled-coil conformations within the HP.

### 2.6. Stability and Predicted Protein Interactions (PPI) assessment

A system called DISULFIND (http://disulfind.dsi.unifi.it/) can calculate the disulfide state and disulfide connectivity of cysteines using just their sequence. Furthermore, many proteins rely on disulfide bridges to stabilize their folding process. In order to explore the structural and functional features of certain proteins, it is crucial to identify disulfide bridges (Ceroni et al. 2006). A database of estimated and known protein interactions, STRING 12.0 (Szklarczyk et al. 2023), is available. The four sources of these interactions include high-throughput tests, conserved co-expression, genomic context, and prior information. They include both direct (physical) and indirect (functional) relationships. In order to convey information across species when necessary, STRING quantitatively combines interaction data from several sources for many organisms.

### 2.7. Secondary and Three-dimensional structure prediction

The PSIPRED and SOPMA servers were used to predict the secondary structure of the HP. PSIPRED (http://bioinf.cs.ucl.ac.uk/psipred/) (McGuffin et al. 2000) is a PSI-blast-based secondary structure prediction tool, while SOPMA (https://npsa.lyon. inserm.fr) (Geourjon and Deléage 1995) is a self-optimized prediction method from an alignment tool. The secondary structural components of putative proteins that have not been characterized were determined using the web-based service I-TASSER (Iterative Threading ASSEmbly Refinement) (Yang and Zhang 2015). One of the bioinformatics tools for making 3D protein models is I-TASSER. The highest Template Modelling (TM) value and C-score were employed to choose the most suitable model from among the five that were developed. The final structure was represented using the PyMol software package (Delano 2002).

### 2.8. Protein Refinement and Model quality assessment

With ModRefiner (http://zhanglab.ccmb.med.umich.edu/ModRefiner/) (Xu and Zhang 2011), protein structures may be built and refined from C-alpha traces. A composite force field, both physics-based and knowledge-based, is used to improve the side chains and backbone atoms once the main-chain structures are generated from the C-alpha traces (Herráez 2006). ModRefiner improves both global and local structural features, resulting in more accurate side-chain positioning, enhanced hydrogen-bonding networks, and reduced atomic clashes.

MolProbity (http://molprobity.biochem.duke.edu/) (Williams et al. 2018), Verify3D (http://nihserver.mbi.ucla.edu/Verify_3D/) (Eisenberg et al. 1997), and the ERRAT Structure Evaluation site (https://servicesn.mbi.ucla.edu/ERRAT/) (Colovos and Yeates 1993) were used to examine the quality of the HP’s predicted three-dimensional structure. The phi and psi bonds of the protein structure are represented in the Ramachandran plot for assessment of the model quality.

### 2.9. Protein Active site detection and Ligand preparation

The protein’s active site was found using an internet-based server called Computed Atlas of Surface Topography of Proteins (CASTp) (http://sts.bioengr.uic.edu/castp/) (Dundas et al. 2006). Detecting ligand-binding sites is a crucial task for the modeled protein, as it is essential for conducting further docking studies. This analysis provides insights needed to create a grid before performing docking experiments, thereby facilitating drug discovery (Ittiyavirah and Paul 2013).

Traditional medicinal plants have been used to extract essential natural bioactive components that serve as significant sources for antimicrobial treatments. An archive of 1,092 chemical compounds extracted from 17 conventionally used medicinal flora was established via the existing GC-MS results through an extensive literature review (**Supplement Table 6**). All compounds, including the control (Nitazoxanide), were retrieved from the PubChem database (https://www.pubchem.ncbi.nlm.nih.gov/) in 2D as well as 3D SDF formats. The PyRx program was used to evaluate the ligand structures. To lower the ligand energies, the gradient descent optimization approach was used in conjunction with the MMFF94 force field. By following this process, the ligands will take on an energetically favorable configuration, rendering them appropriate for subsequent molecular docking investigations.

### 2.10. Molecular Docking

In the drug discovery procedures, molecular docking is a high-throughput analysis to understand the molecular interactions with favorable bonding that are fundamental to key biological functions (Jakhar et al. 2020). This powerful technique offers a comprehensive insight into the interactions between molecules, facilitating the discovery of potential drug candidates (Naqvi et al. 2019). This study involved conducting molecular docking analyses on a library of 1,092 compounds to identify potential inhibitors for our HP. The AutoDock Vina screening program (Cosconati et al. 2010) was utilized for structure-based virtual screening, with all small chemical and receptor structures preprocessed in PDBQT format. A structurally blind search was employed in the virtual screening process, enabling the compounds to navigate and explore their binding sites on the protein. The docking of the compounds was conducted with the grid box’s central location at X: 75.5773, Y: 75.8267, Z: 75.9794. The dimensions were set at X: 78.0801, Y: 45.3813, and Z: 81.0858 Å. A negative binding value was indicated by the ligand’s binding affinity, which was calculated in kcal/mol (Trott and Olson 2010). The binding hits with the lowest energy scores have been identified based on their binding affinities and scoring functions. Additionally, re-docking was performed to verify their binding affinity. Lower energy values suggest a better affinity for the protein binding site. The molecular visualization software PyMOL (Yuan et al. 2017) was utilized to analyze and visualize the docked conformations of each ligand screened against the HP protein. This advanced tool enables the creation of high-quality, comprehensive representations of proteins and chemical substances, providing both three-dimensional and two-dimensional animated visualizations. Beyond its visualization capabilities, PyMOL, ChimeraX (Meng et al. 2023), and Discovery Studio Client software facilitates accurate measurements of various molecular parameters, including bond lengths and the distances between proteins and ligands, offering essential insights into molecular interactions.

### 2.11. Bioactivity and ADME/T properties analysis

Subsequent to molecular docking, the selected compounds undergo further investigation to evaluate their drug-likeness and ADME properties. The ADME (absorption, distribution, metabolism, and elimination) characteristics of two chosen drugs were assessed using the SwissADME and pKCSM web servers. Additionally, a Molecular inspiration Cheminformatics service has been used to ascertain the biological characteristics of the two efficacious chemicals using molecular docking. In this scenario, a chemical with a bioactivity score beyond 0 is deemed biologically active; values ranging from -5.0 to 0 indicate moderate biological activity, whereas scores below -5.0 signify that the molecule is biologically ineffective. The initial evaluation of a potential compound’s cytotoxicity is a crucial phase in pharmaceutical development and research. Conventional toxicity assessment methods involving live animal models are often costly, time-intensive, and raise significant ethical concerns. In order to avoid these restrictions, *in silico* techniques, including computer-assisted development of drugs, provide a more effective and acceptable option for toxicity evaluation. To determine how harmful the chemicals were, this study employed the ProTox-III (Banerjee et al. 2018) server. It was possible to get toxicity estimates for each medication by providing the server with the standard SMILES (simplified molecular-input line-entry system) notation.

### 2.12. MD simulation

One useful tool for studying the behavior and mobility of molecules and atoms over time is MD simulations. In order to analyze the structural changes that occurred during the modelling phase, this work simulated the docked complexes. All simulations conducted utilizing the YASARA dynamics software used the AMBER14 (Krieger and Vriend 2015) force field for computational research. The particle mesh Ewald technique (Krieger et al. 2006) was utilized to determine long-range electrostatic interactions, whereas the examination of short-range van der Waals and Coulomb interactions was conducted with an 8 Å cutoff radius. Pressure and temperature were effectively regulated throughout the simulation using a Monte Carlo barostat and a Langevin thermostat (Mahmud et al. 2021). In order to simulate the human body, the system was controlled at 298 K, a saline content of 0.9% NaCl, and pH 7.4 with water acting as the solvent. The steepest descent method was used to minimize the system (Arshia et al. 2021). To assist trajectory data processing, we conducted the MD simulation at 100 picosecond snapshot intervals and a simulation time step of 1.25 fs (Krieger et al. 2006) for 100 ns. In order to analyze the trajectory data thoroughly, the MD simulation was run at 100 ps snapshot intervals. The simulation of MD relies on the forces acting upon the atoms to iteratively calculate their positions and velocities. For every simulation trajectory, we measured the RG, the hydrogen bond interactions, the solvent accessible surface area, the root mean square deviation, and the root mean square fluctuation (Durrant and McCammon 2011; Khan et al. 2020; Mahmud et al. 2021; Hossain et al. 2023). Using Matplotlib, the MD simulation graphs were generated.

### 2.13. Density functional theory (DFT) calculation

Density Functional Theory (DFT), a quantum physics technique, has the potential to properly portray electron configurations in molecules. Electronic characteristics, structural configurations, and energy levels are among the molecular quantities that may be computed in this way. This quantum mechanical calculation was carried out using the software package Gaussian 09 W. The electronic properties of the molecules were assessed in their singlet ground state, under conditions excluding external energy and solvent effects, utilizing the 6-31G(d) basis set in combination with the Hartree-Fock approach within the framework of DFT. In order to assess molecular reactivity, the DFT method examined several reactivity attributes. Chemical softness (ζ), chemical hardness (η), ionization potential, electronegativity (χ), electrophilicity (ϋ), and electronic potential (μ) were among the features listed (Legler et al. 2015). The DFT approach was employed to analyze various reactivity descriptors, enabling a comprehensive evaluation of the molecular reactivity. The properties encompassed chemical hardness (η), chemical softness (ζ), electron affinity, ionization potential, electronegativity (χ), electronic potential (μ), and electrophilicity (ω). These descriptors were calculated using Koopmans’ theorem, based on the energies of the frontier molecular orbitals (Parr et al. 1999). The following expressions were used:

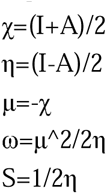

Here, I represent the ionization potential, approximated as the negative of the HOMO energy, and A is the electron affinity, approximated as the negative of the LUMO energy.

### 2.14. Binding free energy calculations using MM-GBSA

The MM-GBSA method was frequently employed to measure the binding free energy of complexes composed of protein-ligand. This approach is highly effective for estimating the binding free energy in molecular complexes, particularly in receptor–ligand interactions. The binding free energy was computed for all top-scoring compounds, the protein, and the known inhibitor. The Prime MMGBSA technique was used to determine many energy attributes, including covalent binding energy, lipophilic energy, Van der Waals energy, Coulomb energy, total energy (Prime energy), generalized Born electrostatic solvation energy, and hydrogen-bonding energy.

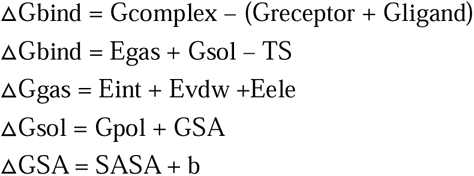

Consequently, the above approaches may provide greater insight into the observed ligand-receptor complex and associated structure.

## 3. Result and Discussion

### 3.1. Characterization of the HP TYC33605.1

#### Multiple sequence alignment and phylogenetic tree construction

The BLASTp analysis against the non-redundant and SwissProt databases indicated similarities of our HP with other bacterial sensor domain-containing DGC proteins. (**Table 1** and **Supplement Table 2**).

**Table 1:** Similar protein obtained from non-redundant UniProt KB/SwissProt sequences.

| Protein ID | Organism | Protein name | Identity (%) | Score | e-value |
| --- | --- | --- | --- | --- | --- |
| WP_000350040.1 | *Vibrio cholerae* | Diguanylate cyclase | 100.00% | 954 | 0.0 |
| AAF96459.1 | *Vibrio cholerae* O1 biovar El Tor str. N16961 | GGDEF family protein | 100.00% | 953 | 0.0 |
| WP_171356651.1 | *Vibrio cholerae* | Diguanylate cyclase | 99.78% | 953 | 0.0 |
| NOE85880.1 | *Vibrio cholerae* | Sensor domain-containing diguanylate cyclase | 99.78% | 952 | 0.0 |
| HAS7850826.1 | *Vibrio cholerae* | TPA: Sensor domain-containing diguanylate cyclase | 99.78% | 952 | 0.0 |
| WP_000350041.1 | *Vibrio cholerae* | Diguanylate cyclase | 99.78% | 952 | 0.0 |
| MVB15607.1 | *Vibrio cholerae* | Sensor domain-containing diguanylate cyclase | 99.78% | 951 | 0.0 |
| EKM9578803.1 | *Vibrio cholerae* | Sensor domain-containing diguanylate cyclase | 99.78% | 951 | 0.0 |
| WP_053032761.1 | *Vibrio cholerae* | diguanylate cyclase | 99.78% | 951 | 0.0 |
| AUR71756.1 | *Vibrio cholerae* | Sensor domain-containing diguanylate cyclase | 99.78% | 951 | 0.0 |

The FASTA sequences of the putative protein (TYC33605.1) were compared to the related annotated proteins by multiple sequence alignment. Phylogenetic analysis was employed to validate the homology evaluation among the proteins, encompassing both individual and subunit levels. Based on BLAST and alignment results that provide a similar understanding of the protein, a phylogenetic tree was built represented in **Supplement Fig 1**.

#### Physiochemical features of the protein

A total of 461 amino acids have been identified in the protein, with the following amino acids being present in the greatest quantity Met (10), Sec (0), Trp (7), His (9), Lys (21), Gln (24), Asp (25), Gly (27), Val (28), Pyl (0), Cys (4), Pro (11), Tyr (15), Arg (29), Glu (31), Asn (19), Thr (21), Ser (33), Ala (40), Leu (58), Ile (24), and Phe (25). The computed molecular weight of the protein is 52,254.72 Daltons, and its theoretical isoelectric point (pI) of 5.93 suggests that the protein carries a net negative charge under physiological conditions. Depending on their amino acid composition, proteins can have isoelectric points (pI) ranging from strongly acidic to highly basic, typically between pH 4.0 and 12.0. When the surrounding pH exceeds the pI, proteins tend to acquire a negative charge; conversely, they become positively charged when the pH falls below the pI. At the pI, the net charge of the protein is neutral, meaning the positive and negative charges are balanced. These pI values have been historically used to differentiate proteins in various techniques such as protein isolation, separation, purification, and crystallization (Tokmakov et al. 2021).

The protein’s molecular formula was determined to be C2348H3677N637O686S14. A detailed overview of its physicochemical properties, such as amino acid composition and distribution, is presented in **Supplementary Fig 2** and **Table 2**. Under in vivo conditions, the protein’s half-life was estimated to exceed 10 hours in Escherichia coli and 20 hours in yeast, while in mammalian reticulocytes (in vitro), it was approximately 30 hours. Additionally, the analysis revealed a higher number of negatively charged residues (Asp + Glu = 56) compared to positively charged ones (Arg + Lys = 50), suggesting a slightly acidic character. The extinction coefficient for HP at 280 nm ranges from 61100 to 60850 M^-1^ cm^-1^ using Expasy’s Protparam. High levels of tryptophan (Trp), tyrosine (Tyr), and cysteine (Cys) indicate a greater extinction coefficient for HPs. The calculated extinction coefficient is applicable for quantitative analyses of protein-ligand and protein-protein complex interactions in solution. The instability index value of HP’s ranges from 30.44 to 50.35. Proteins exhibiting an instability index under 40 tend to be stable, whereas those with a value exceeding 40 are considered unstable (Guruprasad et al. 1990). Another parameter for identifying protein structure is the instability index. The instability index (II) of the target protein was anticipated to be 36.34, categorizing it as stable. This statistic indicates how stable the protein is in a test tube. A protein is deemed unstable if its instability index exceeds 40 and stable if the index is below 40. The Aliphatic Index (AI) measures the fraction of a protein’s volume ascribed to aliphatic side chains (A, V, I, and L), which serves as a thermal stability indicator for proteins with globular structures. The AI value for the HP is 95.66. Proteins exhibiting a high AI demonstrate stability across a broad temperature spectrum, whereas those with a lower AI show reduced thermal stability and increased flexibility. The grand average hydropathy (GRAVY) value of HP is - 0.143. Hydrophilic proteins have negative GRAVY values, while hydrophobic proteins have positive values. Low GRAVY allows for greater protein-water interaction. The GRAVY (Grand Average of Hydropathy) score of a protein is calculated by summing the hydropathy indices of all amino acids in the sequence and dividing the total by the number of residues (Ikai 1980).

**Table 2:** Physicochemical properties of the HP TYC33605.1 predicted using ProtParam.

| Parameters | Values |
| --- | --- |
| Total amino acid residues | 461 |
| Molecular weight | 52254.72 |
| Theoretical isoelectric point (pI) | 5.93 |
| Total count of anionic residues (Asp + Glu) | 56 |
| Total count of positively charged residues (Arg + Lys) | 50 |
| Aliphatic index | 95.66 |
| Instability index (II) | 36.34 |
| GRAVY | -0.143 |
| All Cys residue pairs generate cystines; extinction coefficients | 61100 |
| Extinction coefficients (all Cys residues are reduced) | 60850 |

#### Subcellular localization of protein

CELLO was used to predict subcellular localization, while PSLpred, PSORTb, and SOSUIGramN verified the results. The putative protein was determined to be localized in the inner membrane (**Supplement Table 3**). The existence of transmembrane helices, predicted by HMMTOP and THMM, further validates this finding. The query protein’s status as a transmembrane protein was also verified by the CCTOP service. The protein is categorized as a membrane protein based on all of these findings.

#### Functional annotation of the protein

This HP sequence was shown to possess two unique domains via the conserved domain (CD) search tool, in addition to non-specific domains. The specific domains are PBP2_HisK_like_1 (accession no: cd13706) from amino acids 31 to 251 and GGDEF (accession no: COG2199) from amino acids 191 to 457, both of which belong to the bacterial DGC protein family. The DGC family is known for synthesizing c-di-GMP, a second messenger that has recently been recognized as a key signal in controlling biofilm formation and inhibiting motility (Römling et al. 2005; Jenal and Malone 2006; Ryan et al. 2006b; Cotter and Stibitz 2007; Tamayo et al. 2007). The non-specific domains include SBP_bac_3 (accession no: pfam00497) from amino acids 33 to 250 and GGDEF (accession no: cd01949) from amino acids 304 to 456 (**Supplement Table 4**). The synthesis of c-di-GMP is carried out by DGC enzymes that contain GGDEF domains. On the other hand, phosphodiesterase (PDE) enzymes, which include either an EAL or an HD-GYP domain, are responsible for the breakdown of c-di-GMP (Ryjenkov et al. 2005; Dow et al. 2006; Ryan et al. 2006a). Biofilms have historically been extremely challenging to eliminate with small-molecule drugs. However, uncovering c-di-GMP-mediated mechanisms involved in biofilm formation has opened up the possibility of developing a new generation of anti-biofilm therapeutics that specifically target c-di-GMP signaling pathways (Rabin et al. 2015). Two motifs, including DGC (PF00990) and extracellular solute-binding proteins (PF00497), were identified through Pfam using the motif server (**Supplement Table 5**). Furthermore, the Superfamily search in **Fig 2** indicated the presence of a Bacterial DGC superfamily (accession no. IPR050469). This family produces c-di-GMP, which controls cell surface features such as biofilm formation, cellulose synthesis, and motility. Typically, members contain a GGDEF domain that catalyzes two GTP molecules to create c-di-GMP. Some members also participate in signal transduction pathways and may have additional domains for sensory functions, such as oxygen sensing through a globin-fold domain. The DGC family is crucial for transitioning between different bacterial lifestyles, influencing both environmental persistence and host-pathogen interactions. Fold pattern recognition by the PFP-FunDSeqE tool identified a ‘periplasmic binding protein-like’ fold within the protein sequence. Periplasmic binding proteins (PBPs) function as non-enzymatic receptors in bacteria, facilitating the detection and transport of small molecules into the cytoplasm. They often play a role in the transport of solute molecules through ABC transporters (Tam and Saier 1993). They target vital substances such as vitamins, ions, amino acids, and carbs. PBPs are also involved in QS, chemotaxis, and other signaling systems (Quiocho and Ledvina 1996; Chen et al. 2002). The x-axis in the coiled-coil graph in **Supplement Fig 3** represents the amino acid position in the protein, beginning at the N-terminus, while the y-axis displays the probability score of the coiled-coil. ’Window’ describes the breadth of the ’window’ of amino acids that are scanned at a single moment. The DISULPHIND server indicated the absence of disulfide linkages in these proteins, signifying their heat instability. Disulfide bridges play a critical role in defining the structural and functional properties of numerous proteins and are essential for stabilizing their proper folding during the maturation process.

**Fig 2:**
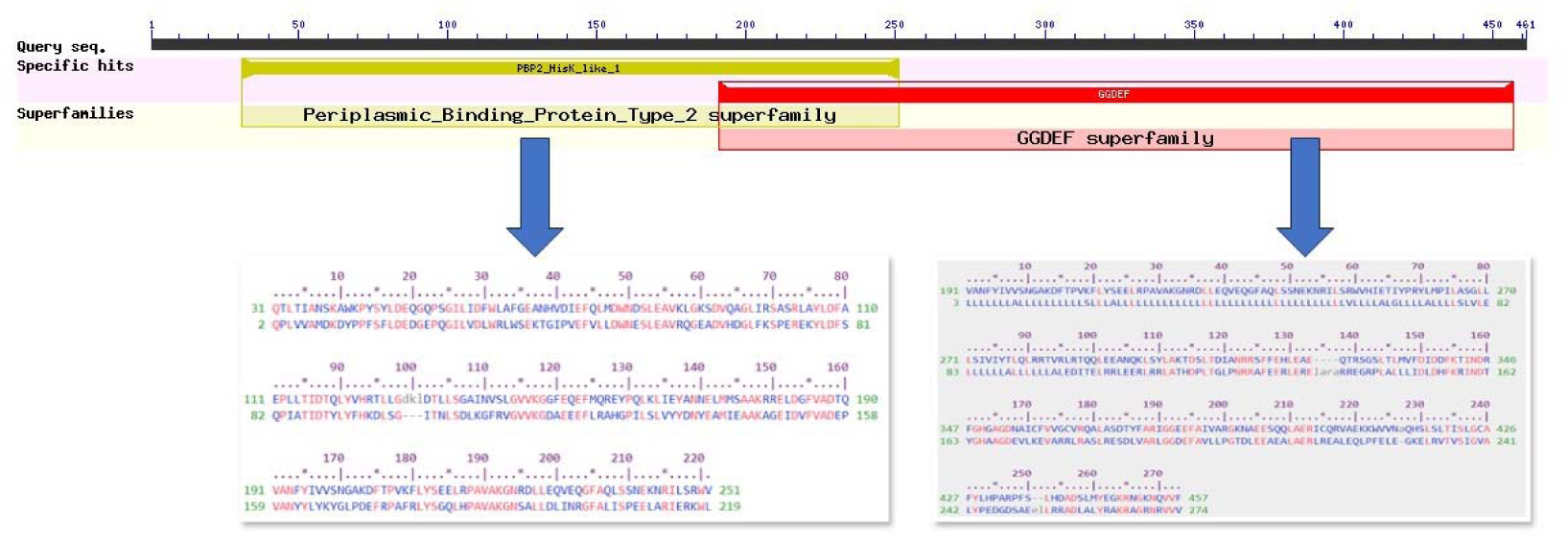
Functional annotation of TYC33605.1 showing domain architecture and conserved motifs. Domains (PBP2_HisK_like_1 and GGDEF) and motifs (GGDEF, SBP_bac_3) are highlighted, supporting roles in signal transduction and biofilm regulation.

#### Secondary and three-dimensional structure analysis

Secondary structure analysis is advancing rapidly to better predict protein function and structure. Using SOPMA with standard parameters, the secondary structures of HPs were predicted. **Supplementary Fig 4** illustrates that the protein’s secondary structure comprises about 53.36% alpha helices, 15.84% extended strands, and 30.80% random coils. This program identified various structural elements: Pi helix (Ii), Alpha helix (Hh), 3_10_ helix (Gg), Bend region (Ss), Extended strand (Ee), Beta bridge (Bb), Beta turn (Tt), Random coil (Cc), and Ambiguous states.

The conventional methods for determining three-dimensional protein structures are NMR spectroscopy and X-ray diffraction. The homology protein structure modeling method was employed to predict 3D homology models for the TYC33605 protein, which represents a substantial advancement in computational biology. Homology modeling is a highly effective method for investigating molecular evolution, as proteins that are evolutionarily related often share a particular structure. The I-TASSER server, a strong tool for automatically guessing protein structure and function, was used to figure out the three-dimensional structure of the TYC33605 protein. The three-dimensional conformation of the TYC33605 protein in **Fig 3** was determined by homology modeling, utilizing template proteins obtained from sequence alignments of published structures in the PDB. The verification of the three-dimensional structure relies on stereochemical parameters, especially the torsion angles that influence protein folding. The restrictive range of these angles means that even minimal variations in torsion angles can provide an accurate estimation of the protein’s correct structural conformation (Tosatto and Battistutta 2007).

**Fig 3:**
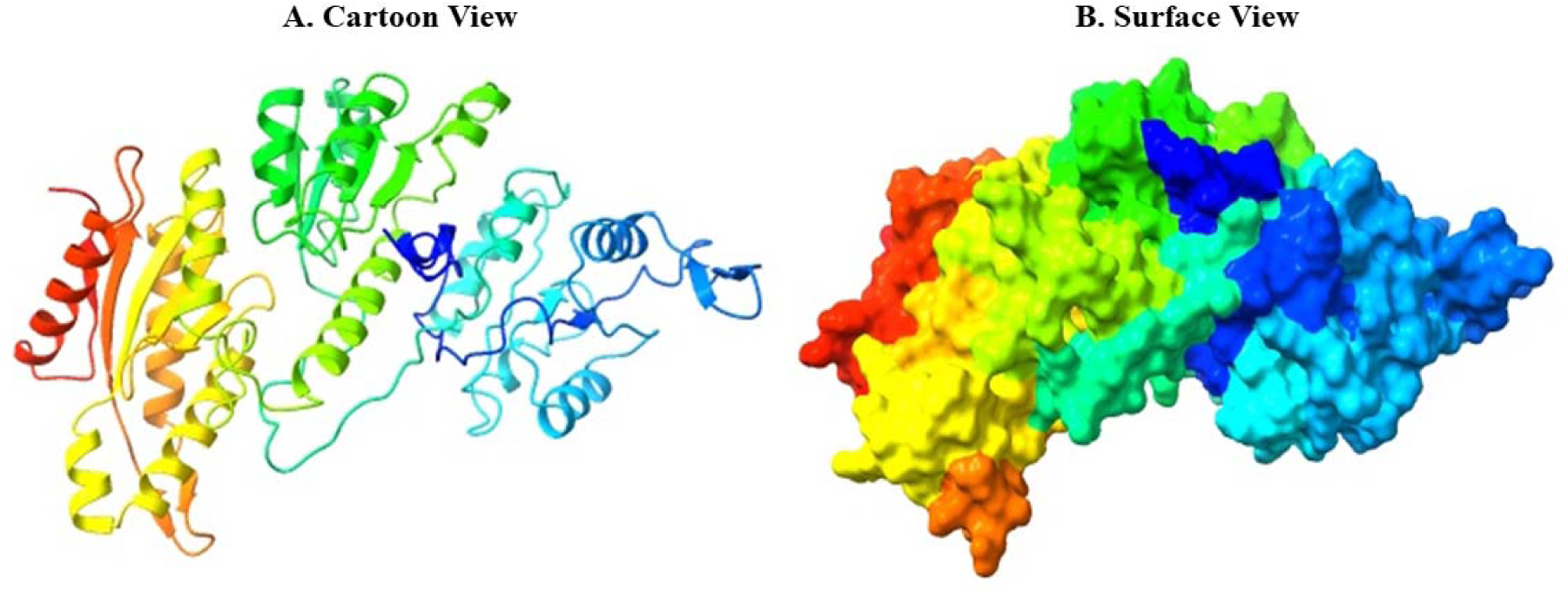
Homology model of TYC33605.1 generated by I-TASSER and visualized in PyMOL and ChimeraX.

The quality of the predicted three-dimensional structure was further confirmed by ERRAT and Verify3D validation tools. ERRAT reported a high overall quality factor of 90.62, while Verify3D analysis showed that 64.43% of the residues achieved a 3D-1D score of ≥ 0.1, reflecting a favorable environmental compatibility of the model. Additionally, structural validation using Ramachandran plot analysis via MolProbity confirmed that 91.1% (418/459) of residues were located in favored regions and 98.9% (454/459) in allowed regions, with only five residues falling in disallowed regions (**Supplementary Fig 5**). These results indicate that the backbone torsion angles (φ and ψ) fall within acceptable limits, supporting the reliability of the model.

#### Protein-protein binding analysis

Protein-protein interactions (PPIs) are essential for almost all cellular functions.

Proteins frequently engage in interdependent interactions to execute a shared function. Translational factors interact with one another to facilitate the entire translation process. A protein’s function can often be inferred based on its interactions with other proteins. Proteins rarely exhibit functionality through interactions with other biomolecules. Gene Ontology (GO) biological processes analysis through the STRING server is represented in **Supplementary Fig 6.**

In this post-genomic era, PPI databases have become one of the most important ways to look for molecular networks and processes in cells (Laurie and Jackson 2005). The recognized functional units with values were VC_A0785 (0.959), VC_1086 (0.958), VC_1710 (0.953), VC_1349 (0.951), VC_0303 (0.944), VC_2453 (0.936), VC_2369 (0.931), VC_A0709 (0.916), VC_A0931 (0.910), and VC_0072 (0.909). Of them, VC_A0785 is a possible membrane protein shown in **Fig 4**.

**Fig 4:**
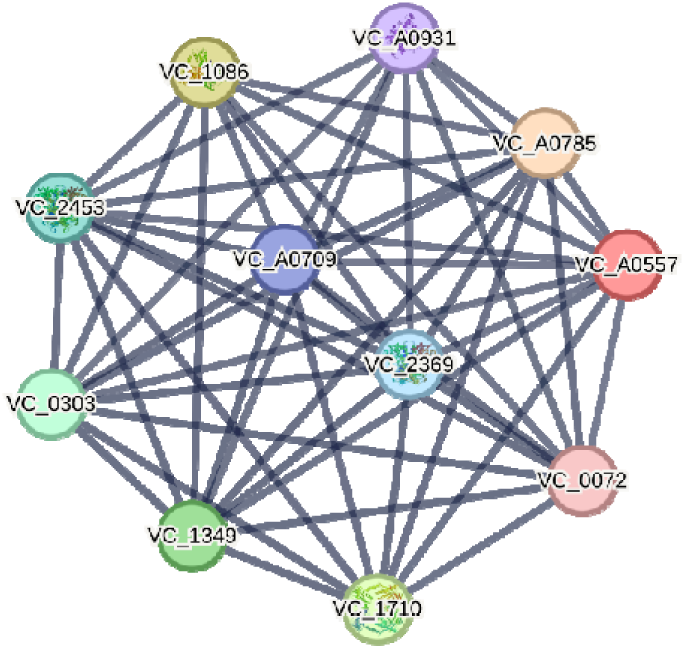
Protein-protein interaction (PPI) network of TYC33605.1 (STRING). Interacting partners (e.g., VC_A0785, VC_1086) are involved in biofilm regulation and c-di-GMP signaling. Edge thickness reflects interaction confidence (scores: 0.9–0.95).

#### MD simulation analysis of the HP

MD simulations are critical for understanding the dynamic behavior of biomolecular structures in solution because they precisely follow their motions over several time frames. MD simulations offer valuable insights into molecular properties, capturing thermal averages that closely align with experimental ensemble measurements (Katiyar and Jha 2018). Through the computation of bulk fluid properties and the evaluation of free energy changes, MD simulation becomes an indispensable tool for unraveling processes like ligand binding (Maria-Solano et al. 2018). The apo structure of the HP was subjected to a 100-nanosecond MD simulation in order to assess its structural stiffness and conformational stability. Key parameters for the HP and the most promising ligand-bound complex were examined to determine structural stability and dynamic behavior. These included RMSD, Rg, SASA, and RMSF.

#### RMSD and RMSF

The protein structural and stability changes are evaluated using the RMSD analysis. Minimal fluctuations suggest a stable atomic backbone, and a lower RMSD value is regarded as a positive indicator of the docked complex’s stability (He et al. 2023). This study evaluated the apo form of the HP to establish its baseline stability in the absence of ligands. The mean Cα RMSD value for the apoprotein was calculated to be 8.55 Å over the simulation period. As depicted in **Fig 5A**, the RMSD trajectory shows that the apoprotein initially undergoes structural adjustment, with the RMSD increasing from 0.724 Å at 0 ns to approximately 7.0 Å by 7.5 ns. This early rise indicates a phase of conformational adaptation. The RMSD subsequently stabilizes within a range of 7.0 Å to 9.5 Å for the remainder of the simulation, although occasional peaks, such as one reaching 10.31 Å at 51.5 ns, are observed. Despite these fluctuations, no significant structural deviations occurred, indicating that the apoprotein retains a relatively stable fold while exhibiting moderate flexibility. Root Mean Square Fluctuation (RMSF) is an essential metric for evaluating the residue-level flexibility of a protein receptor in the context of ligand interactions during simulation. It measures the extent of atomic displacement in relation to their average locations during MD simulations or comparable dynamic investigations (Ghahremanian et al. 2022). In this study, RMSF values for individual residues were calculated over a 100-ns MD simulation, providing insights into the structural stability and mobility of the HP in **Fig 5B**. The average RMSF across all residues was 3.02 Å, which indicates moderate overall flexibility. However, several regions showed pronounced fluctuations, with the highest values ranging from 5 to 8 Å well above the average. The most significant region of increased fluctuation was observed between residues 40 and 50, where multiple residues exhibited RMSF values exceeding 5 Å. Specifically, TRP 41 (5.17 Å), PRO 43 (5.497 Å), PRO 44 (6.154 Å), SER 45 (5.563 Å), TYR 46 (7.49 Å), LEU 47 (8.205 Å), ASP 48 (8.115 Å), GLU 49 (6.583 Å), and GLN 50 (6.941 Å) were the most flexible, with LEU 47 showing the peak fluctuation at 8.205 Å. This region likely represents a loop or solvent-exposed area that may be involved in conformational changes or ligand binding. Another notable region of high fluctuation was found between residues 196 and 205, with ILE 196 (6.052 Å), VAL 197 (5.655 Å), ASN 200 (5.75 Å), ALA 202 (5.651 Å), LYS 203 (7.294 Å), ASP 204 (6.109 Å), and PHE 205 (5.056 Å) displaying elevated RMSF values peaking at LYS 203 with 7.294 Å. These values suggest that this segment plays a key role in the protein’s dynamic behavior, possibly indicating a flexible domain or interaction site.

**Fig 5:**
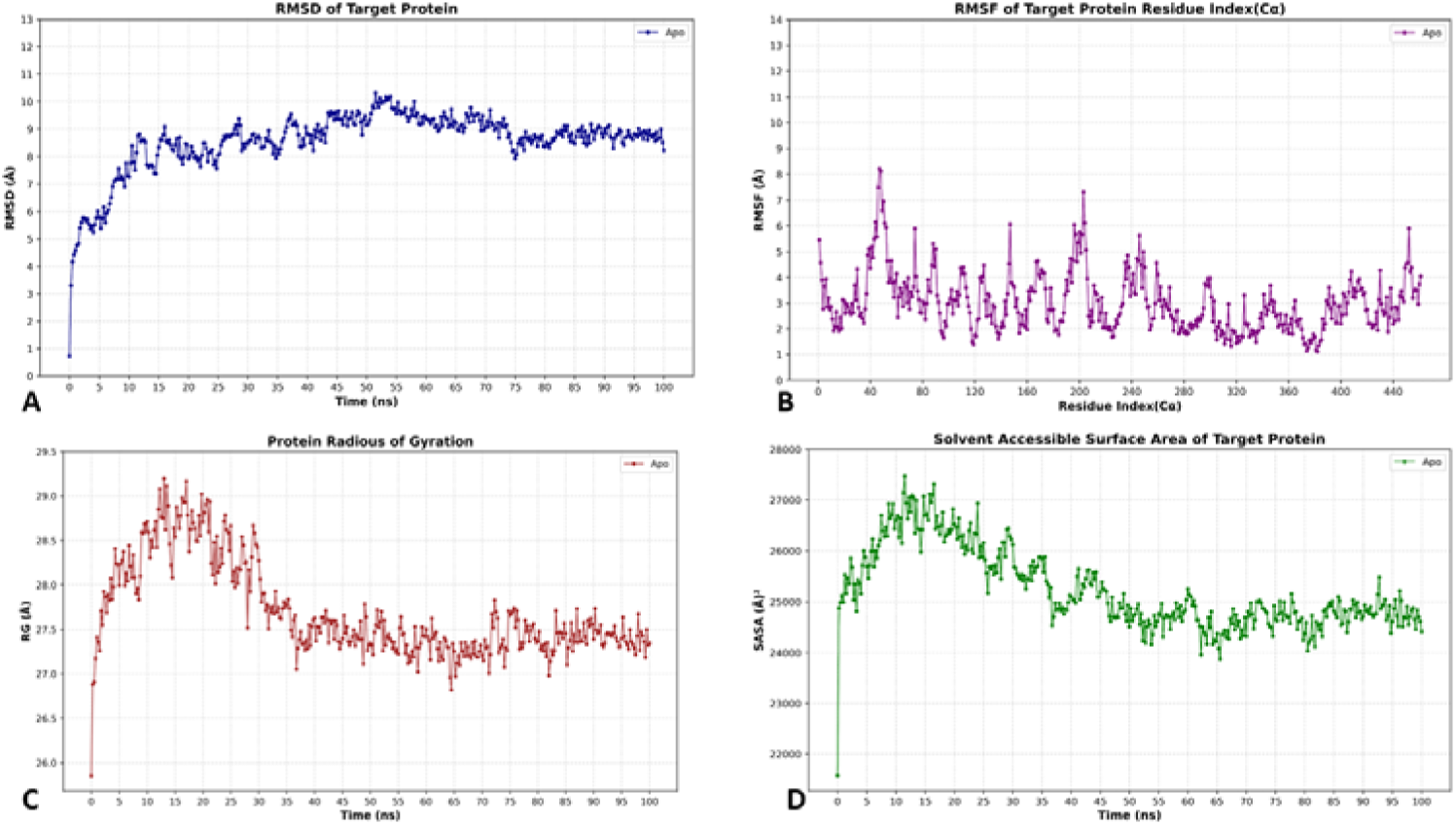
Structural stability and flexibility analysis of the HP TYC33605.1 during a 100 ns MD simulation. RMSD analysis shows the apo protein stabilizes at ∼8.55 Å after initial conformational adjustments (0–10 ns), with a transient peak at 51.5 ns (10.31 Å). RMSF highlights regions of high flexibility, notably residues 40–50 (e.g., LEU47: 8.2 Å) and 196–205 (e.g., LYS203: 7.29 Å), likely in solvent-exposed loops or functional sites. The average RMSF of 3.02 Å indicates stable global folding with localized fluctuations. Radius of gyration (Rg) averages 27.65 Å, with initial expansion followed by stabilization (∼27.3 Å at 100 ns), reflecting a compact structure. Solvent-accessible surface area (SASA) averages 25,227 Å², with peaks (e.g., 27,465 Å² at 11.5 ns) suggesting transient surface exposure during conformational changes. Rg and SASA trends align with structural dynamics revealed by RMSD and RMSF.

#### Rg and SASA

The radius of gyration (Rg) shows how tightly a protein is packed by assessing the spatial distribution of atoms around its central axis over a specified simulation period. This measure tells us a lot about the complex’s general stability and structural integrity along the route (Shrivastava et al. 2022). It also reflects changes in the system’s compactness or expansion, further reinforcing the overall structural conformation and stability. The HP’s Rg values across a 100-ns simulation period are also calculated. In this study, the apo form of the HP was examined to determine its baseline compactness in the absence of ligand interactions. The average radius of gyration (Rg) was recorded at 27.65 Å throughout the simulation period. As depicted in **Fig 5C**, the Rg trajectory initially measured 25.849 Å at 0 ns and progressively increased to around 28.0 Å by 10 ns, indicating an initial structural expansion. Beyond the 10 ns mark, the Rg fluctuated between 27.0 Å and 29.0 Å, with occasional peaks reaching 29.2 Å at 13 ns. Toward the end of the simulation, the Rg stabilized at approximately 27.3 Å, indicating that the apo protein sustains a rather stable degree of compactness with minor structural variations. These findings imply that the unbound protein exhibits moderate flexibility without undergoing significant expansion or compaction.

The area of a protein’s surface that is easily exposed to solvent molecules is measured by its solvent-accessible surface area, or SASA. As demonstrated in **Fig 5D**, it offers molecular insights into the dynamic interactions between protein macromolecules and ligands, as well as conformational fluctuations. This study focused on measuring the protein’s hydrophilic and hydrophobic residues, with a higher SASA score suggesting increased surface exposure. For the apo form of the HP, the average SASA was calculated to be 25,227.35 Å² over the simulation period. As depicted in Fig. 7, the SASA trajectory starts at 21,574 Å² at 0 ns, increases to a peak of 27,465 Å² at 11.5 ns, and concludes at 24,414 Å² by 100 ns. Throughout the simulation, the SASA fluctuates significantly, ranging from a low of 21,574 Å² at the start to the noted peak, with other notable variations such as 23,880 Å² at 65.5 ns. This broad range of SASA values indicates substantial conformational changes and variations in solvent exposure, suggesting that the apoprotein undergoes dynamic structural adjustments while maintaining an overall moderate level of solvent accessibility.

### 3.2 Discovery and Validation of Potential Natural Inhibitors

#### Prediction of the protein active site

Determining the positions of ligand-binding sites on proteins is crucial for several applications, including structural characterization, molecular docking, functional site comparison and de novo drug development. The anticipated active region of the hypothesized protein comprises 42 amino acids important to its high activity. The residues Asn169, Glu171, Leu172, Ala176, Leu247, Trp250, Arg285, Glu293, Ala294, Ala311, Arg364, Leu367, Ser369, and Phe373 were found to be involved in the active site of the HP of *V. cholerae*, confirmed by the CASTp online server. **Supplementary Fig 7** depicts the interaction residues discovered in the active site of TYC33605.1. Finding binding interactions and conducting docking experiments with certain ligands is made easier with the use of this information on active binding site residues.

#### Frontier molecular orbital study (Homo-Lumo)

In quantum chemistry, Frontier Molecular Orbital (FMO) analysis is a method employed to investigate the geometries, sources of energy, and distributions of electrons associated with a molecule’s lowest unoccupied molecular orbital (LUMO) and highest occupied molecular orbital (HOMO).

Molecular orbital behavior, particularly in organic molecules, may be better understood using this approach. The reactivity of molecular orbitals was assessed by analyzing the HOMO-LUMO levels, and it has been shown that molecules with a minimal or insignificant HOMO-LUMO gap generally display increased activity potential. Possible inhibitors were the focus of this investigation. The electron affinity and ionization potential of Nitazoxanide, Sativanone, and Luteolin were evaluated using FMO calculations. Key electronic properties, including ionization potential, hardness, E_HOMO_, electronegativity, electrophilicity, E_LUMO_, chemical potential, ΔEgap, softness, electronic potential, and electron affinity, were included in the computations. The outcomes are shown in **Fig 6**. One of the most important metrics in quantum chemistry for understanding molecular behavior is electronic energy, abbreviated as Eh. Changing Eh means that the security and bonding links between molecules are changing, while higher energy levels mean that the molecules are reacting more. Among the analyzed compounds, Sativanone had the greatest dipole moment (4.9015 Debye), signifying substantial polarity, while Luteolin displayed the lowest dipole moment (3.6220 Debye), suggesting comparatively reduced polarity compared to the control molecule Nitazoxanide. The disparities in charge distribution and chemical polarity were emphasized by the variations in dipole moments. Sativanone exhibited a high degree of polarity, while Luteolin exhibited a significantly reduced degree of polarity. The determined frontier molecular orbital (FMO) energies indicate unique electronic characteristics for each compound. The E_HOMO_ and E_LUMO_ values for Nitazoxanide are -0.373 au and 0.054 au, respectively, which leads to an energy gap (ΔEgap) of 0.427 au (**Fig 6A**). Sativanone exhibits an E_HOMO_ of -0.316 au and an E_LUMO_ of 0.093 au, resulting in a ΔEgap of 0.409 au (**Fig 6B**). In the meantime, Luteolin exhibits an E_HOMO_ of -0.318 au and an E_LUMO_ of 0.053 au, resulting in a ΔEgap of 0.371 au (**Fig 6C**). The energy gap values indicate the electronic stability and potential reactivity of the compounds.

**Fig 6:**
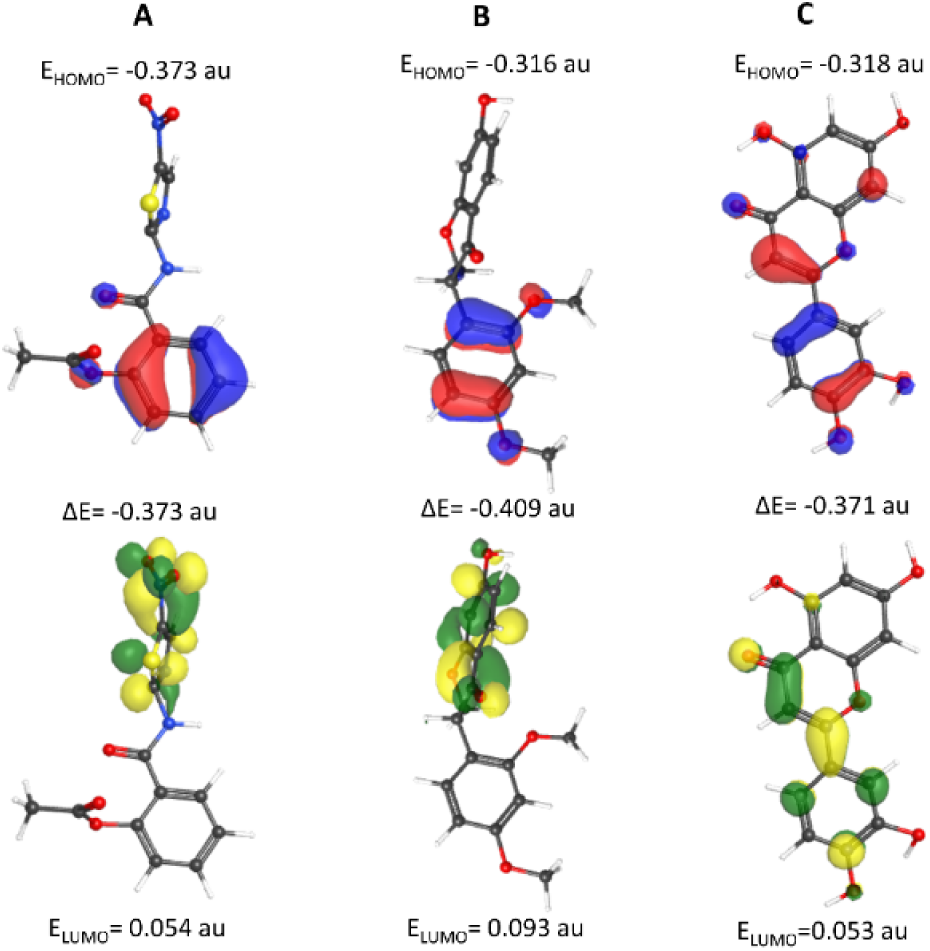
Frontier Molecular Orbital (FMO) analysis of Luteolin and Sativanone. HOMO-LUMO gaps (Luteolin: 4.2 eV; Sativanone: 3.8 eV) indicate electronic stability. Red/blue regions represent electron-rich/deficient areas.

The electronic descriptors reveal significant differences in the electronic structures of the lead compounds compared to the control. Sativanone displays distinct electronic characteristics, marked by a lower E_HOMO_ value and a wider energy gap, suggesting greater stability and reduced electron-donating ability. In contrast, Luteolin exhibits a narrower energy gap and a higher E_HOMO_, indicating enhanced ionization potential and reactivity. Moreover, significant disparities in electronegativity and electrophilicity between the two compounds indicate variations in their chemical reactivity and possible molecular interaction characteristics.

#### Molecular Docking Assessment

TYC33605.1, a putative protein, possesses many essential sites that facilitate the ligand-binding pocket. These residues, Met174, Ser175, Ala176, Lys222, Gly223, Asn224, Arg225, Ile246, Leu247, Trp250, Arg285, Leu286, Gln290, Glu293, Lys303, Asp309, Ile310, Ala311, Ser315, Phe316, Glu318, Arg364, Ser369, Thr371, Tyr372, and Phe373, are all of which are important in inhibiting the function of the protein.

In structural biology and structure-based medication development, molecular docking was employed to assess the binding energies of phytochemicals to the target protein (Hori and Kim 2019). Luteolin (**Supplementary Fig 8A**) and Sativanone **(Supplementary Fig 8B)** were chosen as significant compounds because of their inhibitory activity, favorable docking scores, and broad molecular relationships with the proposed protein. The binding analysis for Luteolin and Sativanone reveals that its extensive network of hydrogen bonding and hydrophobic interactions with the main active site residues suggests that these compounds may effectively disrupt the protein’s catalytic core (**Fig 7**). Luteolin is a flavonoid present in many fruits and vegetables, including celery, broccoli, and onions. It demonstrates a broad spectrum of bioactivities, encompassing antioxidant, anti-inflammatory, anti-cancer, and neuroprotective effects. Luteolin’s antioxidant capacity is attributed to its unique molecular structure, which enables it to mitigate oxidative stress and scavenge reactive oxygen species (ROS), a contributing factor in numerous diseases like cancer and neurodegenerative disorders (Taheri et al. 2021). In **Table 3** we can see that it also exhibited a docking score of -9.1 kcal/mol, indicating a significant binding affinity by forming multiple stabilizing interactions within the active site, including conventional hydrogen bonds with residues such as Glu293 (2.59 Å), Arg364 (2.69 Å), Leu367 (2.52 Å), Thr371 (2.41 Å), and Phe373 (2.25 Å), as well as hydrophobic contacts like Pi-Alkyl interactions with Ala176 (5.33 Å) and Met174 (5.28 Å) and additional Pi-Sigma and Pi-Cation interactions with Ser369 (3.92 Å) and Arg225 (4.88 Å).

**Fig 7:**
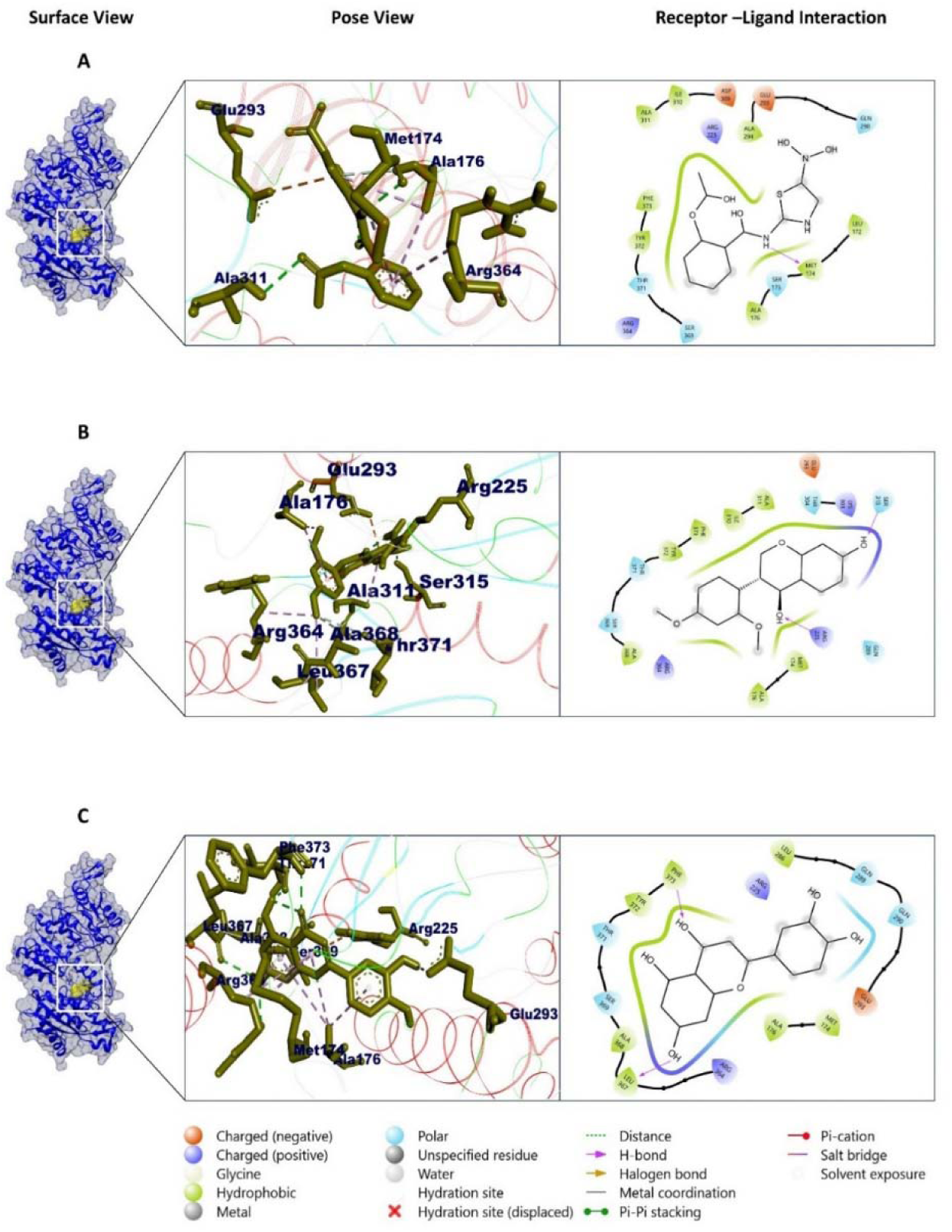
Molecular docking analysis of compounds C0 (Nitazoxanide, control), C1 (Luteolin), and C2 (Sativanone) with the receptor, illustrating surface views, ligand-binding poses, and critical interactions for each compound. Panel A (C0) highlights interactions with Met174, Ala176, Ala311, and Arg364 through hydrophobic contacts and H-bonds, while Panel B (C1) shows engagement with Glu293, Arg225, Ser315, Arg364, and Ala368 via H-bonds, salt bridges (Arg225–Glu293), and π-cation interactions. Panel C (C2) reveals binding to Asp373 and Glu293, involving charged/polar interactions, metal coordination, and displaced hydration sites. Interaction types are denoted by symbols/colors: H-bonds (dashed lines), salt bridges (red), π-cation (orange), metal coordination (blue), and hydrophobic residues (white). Abbreviations include Inr371 (unspecified residue) and Gu367 (potential typo for a metal-coordinating residue). C0 (41884) is the control compound, whereas C1 (5280445) and C2 (13886678) are experimental candidates.

**Table 3:** Molecular docking results of top phytochemicals against TYC33605.1, including binding affinities and interactions.

| Compound name and CID | Predicted pIC50 Score | Binding affinity (kcal/mol) | Residue in contact | Interaction type | Bond distance in Å |
| --- | --- | --- | --- | --- | --- |
| 5280445<br>(Luteolin) | 6.67 | -9.1 | Glu293<br>Arg364<br>Leu367<br>Thr371<br>Phe373<br>Ala176<br>Met174<br>Arg364<br>Ser369<br>Arg225 | H- bond<br>H-bond<br>H-bond<br>H-bond<br>H-bond<br>Pi-Alkyl<br>Pi-Alkyl<br>Pi-Alkyl<br>Pi-Sigma<br>Pi-Cation | 2.587<br>2.685<br>2.516<br>2.414<br>2.245<br>5.329<br>5.276<br>5.213<br>3.923<br>4.881 |
| 6441061<br>(Licarin B) | 6.60 | -9 | Arg285<br>Gln289<br>Glu293<br>Ala311<br>Leu247<br>Lys303<br>Ala311<br>Tyr372 | H-bond<br>H-bond<br>Pi-Anion<br>Alkyl<br>Alkyl<br>Pi-Alkyl<br>Pi-Alkyl<br>Pi-Alkyl | 2.15008<br>2.203<br>3.890<br>3.995<br>5.168<br>4.342<br>4.620<br>4.755 |
| 5280863<br>(Kaempferol) | 6.53 | -8.9 | Glu293<br>Met174<br>Arg364 | Pi-Anion<br>Pi-Alkyl<br>Pi-Alkyl | 3.976<br>5.063<br>5.147 |
| 114829<br>(beta.<br>Methoxybutyraldehyde ) | 6.53 | -8.9 | Leu286<br>Leu367<br>Phe373<br>Glu293<br>Met174 | H-bond<br>H-bond<br>H-bond<br>Pi-Anion<br>Pi-Alkyl | 2.597<br>2.555<br>2.883<br>4.849<br>5.415 |
|  |  |  | Arg364<br>Ala176<br>Leu286 | Pi-Alkyl<br>Pi-Alkyl<br>Pi-Alkyl | 4.068<br>4.018<br>5.469 |
| 128861<br>(Cyanidin) | 6.45 | -8.8 | Ala368<br>Thr371<br>Leu286<br>Arg225<br>Glu293<br>Ala176<br>Arg364 | H-bond<br>H-bond<br>H-bond<br>H-bond<br>Pi-Anion<br>Pi-Sigma<br>Pi-Alkyl | 2.494<br>2.191<br>2.967<br>2.890<br>4.185<br>3.756<br>4.890 |
| 13886678<br>(Sativanone) | 6.38 | -8.7 | Arg225<br>Ser315<br>Ala368<br>Thr371<br>Arg225<br>Glu293<br>Ala176<br>Arg364<br>Leu367<br>Ala311<br>Ala176 | H-bond<br>H-bond<br>C-H bond<br>C-H bond<br>Pi-Cation<br>Pi-Anion<br>Alkyl<br>Alkyl<br>Alkyl<br>Pi-Alkyl<br>Pi-Alkyl | 2.085<br>2.053<br>3.547<br>3.718<br>4.641<br>3.417<br>3.472<br>4.155<br>4.993<br>4.469<br>4.802 |
| 5280961<br>(Genistein) | 6.23 | -8.5 | Arg364<br>Leu367<br>Lys303<br>Arg225<br>Gln289<br>Ala311<br>Arg364<br>Ala311 | H-bond<br>H-bond<br>H-bond<br>H-bond<br>H-bond<br>H-bond<br>Pi-Alkyl<br>Pi-Alkyl | 2.090<br>2.769<br>3.024<br>2.044<br>2.584<br>2.768<br>4.344<br>4.939 |
| 41632<br>(Mepronil) | 6.09 | -8.3 | Met174<br>Arg225<br>Ala176<br>Met174<br>Ala176<br>Met174<br>Arg364 | H-bond<br>H-bond<br>Pi-Donor<br>Alkyl<br>Pi-Alkyl<br>Pi-Alkyl<br>Pi-Alkyl | 2.674<br>2.311<br>3.136<br>4.937<br>4.334<br>5.445<br>4.370 |
| 41684 (Control)<br>Nitazoxanide | 5.57 | -7.6 | Met174<br>ALA311<br>ILE310<br>GLU293<br>ALA176<br>ALA176<br>MET174<br>ALA176<br>ARG364 | H-bond<br>H-bond<br>C-H bond<br>Pi-Anion<br>Pi-Donor<br>Pi-Alkyl<br>Pi-Alkyl<br>Pi-Alkyl<br>Pi-Alkyl | 2.423<br>2.988<br>2.441<br>4.242<br>3.175<br>5.124<br>5.213<br>5.109<br>4.621 |

Robust hydrogen bonding between the ligand and essential protein residues serves as a strong marker for elevated binding affinity, a critical determinant of protein activity. Sativanone, a prenylated isoflavanone predominantly isolated from species within the Dalbergia genus, exhibits a range of bioactivities, including antibacterial and α-glucosidase inhibitory properties. It has demonstrated significant antibacterial activity against Ralstonia solanacearum, a pathogenic bacterium affecting various crops (Subbiah et al. 2024), and acts as a potent α-glucosidase inhibitor with an ECLL of 0.357 mg/mL for rat α-glucosidase, suggesting potential applications in managing conditions like diabetes (Van Bon et al. 2018). In docking studies, Sativanone achieved a score of –9.0 kcal/mol and formed a conventional hydrogen bond with Arg285 (2.15 Å), engaged Gln289 (2.20 Å), established a prominent Pi-Anion interaction with Glu293 (3.89 Å), and participated in hydrophobic contacts with residues including Ala311 (3.99 Å) and Lys303 (4.34 Å). These interactions collectively indicate that Sativanone can form robust binding contacts with the protein’s active site, potentially interfering with its functional dynamics.

#### ADME/Tox and bioactivity analysis

ADME/T analysis assesses a compound’s absorption, distribution, metabolism, excretion, and toxicity, providing a comprehensive evaluation of its pharmacokinetic and toxicological properties. This analysis is essential for determining the compound’s potential as a drug candidate by examining its absorption efficiency, distribution within biological systems, metabolic stability, excretion pathways, and possible toxic effects. Assessing safety concerns, including toxicity and adverse drug reactions, is a fundamental component of drug development. While clinical trials remain essential, computational predictive models for toxicity screening have significantly improved the drug design process by enabling early risk assessment with high accuracy. Evaluating the toxicity and pharmacokinetic properties of lead compounds in **Table 4**, is crucial for determining their efficacy and safety. Pharmacological profiles of the top two potential candidates were derived from the SwissADME server.

**Table 4:** ADME/Tox and pharmacokinetic properties of Luteolin and Sativanone predicted using SwissADME.

| Types of Properties | Properties | Luteolin | Sativanone |
| --- | --- | --- | --- |
| Physico-chemical properties | MW (g/mol) | 286.24 | 300.31 |
|  | Heavy atoms | 21 | 22 |
|  | Aro. Atoms | 16 | 12 |
|  | Rotatable bonds | 1 | 3 |
|  | H-bond acceptors | 6 | 5 |
|  | H-bond donors | 4 | 1 |
|  | TPSA (Å<sup>2</sup>) | 111.13 | 64.99 |
| Lipophilicity | Log Po/w (Cons.) | 1.73 | 2.46 |
| Water Solubility | Log S (ESOL) | -3.71 | -3.58 |
| PK | GI absorption | High | High |
|  | BBB permeant | No | Yes |
|  | P-gp substrate | No | No |
| Drug likeness | Lipinski | Yes, 0 violation | Yes, 0 violation |
| Medi. Chemistry | Synth. accessibility | 3.02 | 3.38 |

In this study, two lead compounds underwent further analysis to assess their pharmacological activity, which plays a vital role in predicting their behavior in the human body. This evaluation was conducted using ADME/T technology, a widely reviewed and increasingly utilized approach in modern drug discovery. Luteolin and Sativanone were evaluated for hepatotoxicity, Central Nervous System (CNS) permeability, Cytochrome P450 (CYP) inhibition, and p-glycoprotein inhibition, among other characteristics (**Table 5**).

**Table 5:** The two selected lead compounds were evaluated for their various toxicity properties.

| Classifications & Endpoints | Target | Luteolin<br>(Class 5) | Sativanone<br>(Class 5) |
| --- | --- | --- | --- |
| Organ Toxicity | Hepatotoxicity | No | No |
|  | Carcinogenicity | Active | No |
|  | Immunotoxicity | No | Active |
|  | Mutagenicity | Active | No |
|  | Cytotoxicity | No | No |
| Toxicity | LD50 (mgkg<sup>-1</sup>) | 3919 | 2500 |

CNS permeability denotes a drug’s capacity to cross the selectively permeable blood-brain barrier, which functions to safeguard the central nervous system from potentially harmful agents. A compound’s ability to successfully pass the blood-brain barrier is typically indicated by a CNS permeability score larger than -4. The molecular weights of Luteolin and Sativanone were 286.24 and 300.31 g/mol, respectively. Both Luteolin and Sativanone conform to Lipinski and Ghosh’s rule of drug-likeness. Luteolin demonstrates superior results in physicochemical properties, water solubility, lipophilicity, PD and PK compared to Sativanone.

#### MD simulation of the protein-ligand complex

MD simulations play a vital role in understanding the dynamic nature of biomolecular structures in solution. By precisely tracking their movements across various timescales. MD simulations offer valuable insights into molecular properties, capturing thermal averages that closely align with experimental ensemble measurements (Saif et al. 2025). Through the computation of bulk fluid properties and the evaluation of free energy changes, MD simulation becomes an indispensable tool for unraveling processes like ligand binding (Katiyar and Jha 2018). Key parameters such as solvent-accessible surface area (SASA), radius of gyration (RG), root-mean-square deviation (RMSD), and root-mean-square fluctuation (RMSF) were analyzed for both the apoprotein and the most promising ligand-bound complex.

#### RMSD and RMSF

The mean Cα RMSD values for CID 13886678 and CID 5280445 in complex with the HP were 8.13 Å and 7.78 Å, respectively. The apo form of the protein exhibited a higher average RMSD of 8.55 Å, while the control compound CID 41684 showed an RMSD of 8.33 Å. CID 5280445 showed the lowest total RMSD of the two compounds, indicating that it maintained the most stable conformation during simulation. This compound demonstrated negligible variations, suggesting a low level of conformational flexibility. These findings show that ligand-bound complexes had smaller fluctuations than the apo form, indicating that ligand interaction helps to maintain the protein’s structural stability over the duration of the simulation. As shown in **Fig 8A**, the overall RMSD profile revealed no major structural shifts.

**Fig 8:**
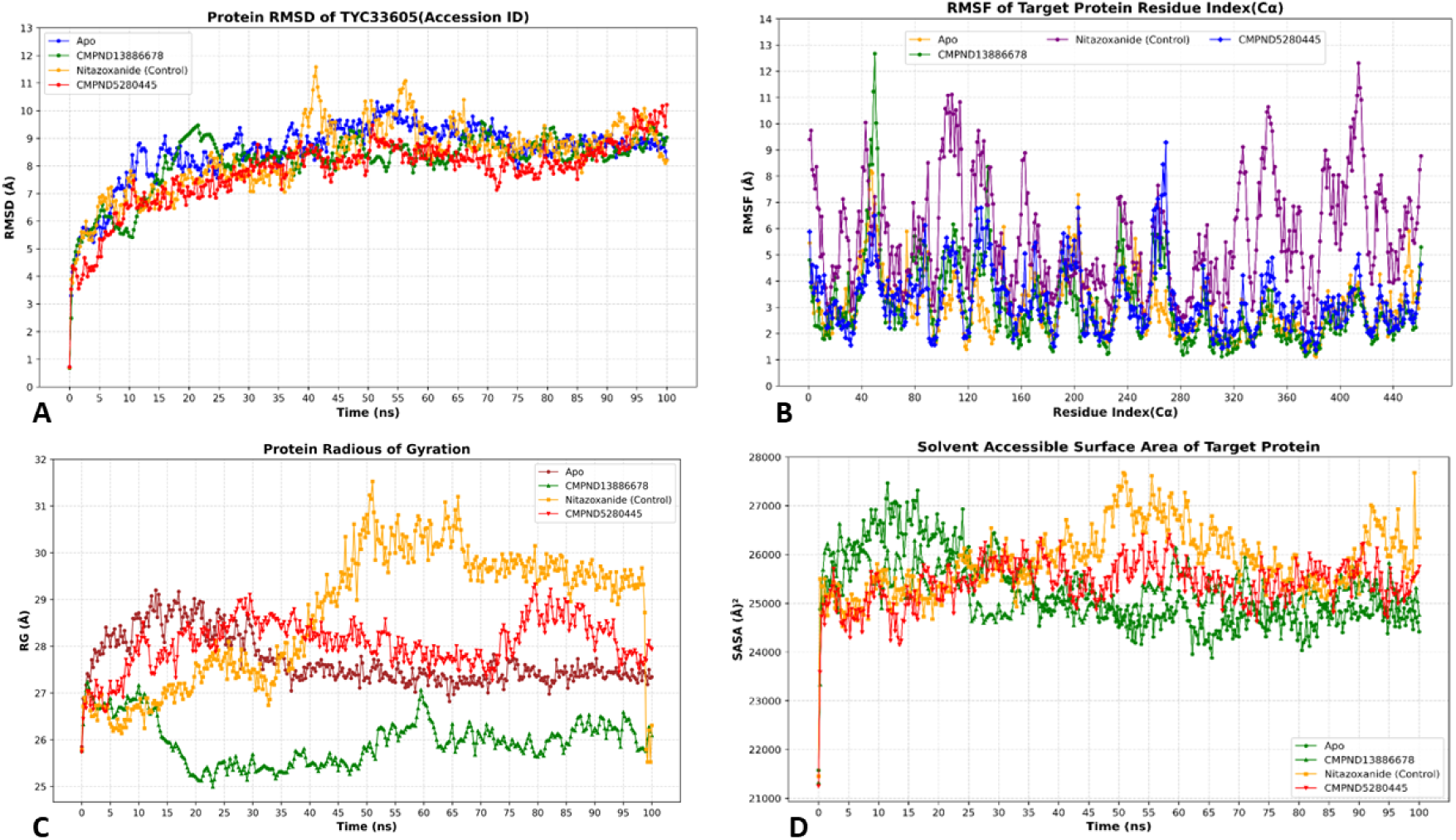
Stability, flexibility, compactness, and solvent accessibility analysis of TYC33605.1 complexes with Luteolin (CID 5280445), Sativanone (CID 13886678), and Nitazoxanide (control) during 100 ns MD simulations. (A) Root Mean Square Deviation (RMSD, Å) tracks backbone stability over time. Sativanone (blue) shows the lowest average deviation, indicating enhanced structural stability compared to Luteolin (green) and the control (red). (B) Root Mean Square Fluctuation (RMSF, Å) maps residue-specific flexibility across residues 1–440. Key dynamic regions (e.g., residues 40–50 and 260–280) are implicated in ligand interactions. Sativanone stabilizes these regions most effectively, while the control exhibits broader mobility, highlighting ligand-specific effects on local dynamics. (C) Radius of Gyration (Rg, Å) measures structural compactness. Sativanone maintains the lowest average Rg (25.97 Å), reflecting a more compact fold than Luteolin (27.91 Å) and the control (27.84 Å). (D) Solvent-Accessible Surface Area (SASA, Å²) indicates solvent exposure. Sativanone minimizes surface accessibility (average SASA: 25,234 Å²), suggesting increased structural rigidity, while the control shows the highest exposure (25,875 Å²). Luteolin exhibits intermediate behavior across all metrics, consistent with moderate binding affinity and stability.

The average RMSF values were 3.03 Å for CID 13886678, 3.25 Å for CID 5280445, and 5.68 Å for CID 41684 (control). Compared to the apoprotein’s average RMSF of 3.02 Å, the complexes with CID 13886678 and CID 5280445 showed slightly higher average fluctuations (3.03 Å and 3.25 Å, respectively), suggesting a minor increase in overall flexibility (**Fig 8B**). In contrast, the CID 41684 complex exhibited a significantly higher average RMSF of 5.68 Å, indicating that this ligand may destabilize the protein structure and lead to increased residue fluctuations. In the CID 13886678 complex, the highest fluctuation was observed at GLN 50 (12.676 Å) with additional elevated values at GLU 49 (11.228 Å), GLY 51 (10.027 Å), LEU 47 (9.398 Å), and GLN 52 (9.06 Å). These values far exceed the complex’s average RMSF of 3.03 Å and suggest that the region around residues 47–52 remains highly flexible, possibly due to its role in the ligand binding site. The CID 5280445 complex showed its peak fluctuation at LEU 269 (9.282 Å) with notable values at SER 267 (8.444 Å), GLY 268 (7.311 Å), ALA 266 (7.259 Å), and PRO 263 (7.132 Å), all significantly above the average RMSF of 3.25 Å. This indicates a distinct flexible region around residues 263–269 that may be influenced by this ligand. The CID 41684 complex displayed the highest fluctuation at GLN 414 (12.315 Å) with significant values at ALA 413 (11.077 Å), ASP 108 (11.115 Å), ALA 105 (11.082 Å), and HIS 415 (11.371 Å). These values are well above the complex’s average RMSF of 5.68 Å and suggest that residues around 105–108 and 413–415 are particularly dynamic in the presence of this ligand, despite the already elevated baseline flexibility. Comparative analysis showed that ligand binding has a varied impact on the protein’s flexibility. While CID 13886678 and CID 5280445 slightly increase the average RMSF compared to the apo form (3.03 Å and 3.25 Å versus 3.02 Å), CID 41684 significantly increases overall fluctuations (5.68 Å), suggesting a destabilizing effect.

#### Rg and SASA

The phytochemicals CID 13886678 and CID 5280445 showed average Rg values of 25.97 Å and 27.91 Å, respectively, when bound to the protein (**Fig 8C**). The control compound, CID 41684, showed a slightly higher average Rg value of 27.84 Å in its complex with the HP, while the apoprotein recorded an average Rg of 27.65 Å. These results suggest that CID 13886678, in particular, maintained a more compact protein structure compared to the control and the unbound form, with CID 5280445 showing values closer to those of the apoprotein and the control. Overall, the relatively stable Rg profiles of CID 13886678 and CID 5280445, particularly the former, suggest that these ligands maintain the protein’s compactness with no substantial structural shifts, in contrast to the more dynamic behavior observed with control compound CID 41684. These findings highlight CID 13886678 as the ligand that most effectively preserves a compact and stable protein structure, which is often a desirable trait for a potential inhibitor. The complex with CID 13886678 exhibited an average SASA of 25,234.28 Å², starting at 21,323 Å² at 0 ns, reaching a peak of 26,630 Å² at 3.5 ns, and concluding at 24,759 Å² by 100 ns. This relatively stable SASA trajectory, as shown in **Fig 8D**, fluctuates between approximately 24,500 Å² and 26,630 Å², indicating minimal changes in solvent exposure throughout the simulation. Similarly, the CID 5280445 complex displayed an average SASA of 25,411.57 Å², beginning at 21,249 Å², peaking at 26,323 Å² at 37 ns, and ending at 25,751 Å². Its SASA profile shows moderate fluctuations, ranging from 21,249 Å² to the peak, with values generally stabilizing around 25,000–26,000 Å², suggesting a consistent level of solvent exposure. In contrast, the control compound CID 41684 exhibited a higher average SASA of 25,875.23 Å², starting at 21,452 Å², peaking at 27,675 Å² at 50.75 ns, and concluding at 26,342 Å². The SASA trajectory for CID 41684 shows greater variability, fluctuating between 21,452 Å² and the peak, with other notable values such as 27,219 Å² at 49.25 ns, indicating increased solvent accessibility compared to the other complexes. When compared to the apoprotein’s average SASA of 25,227.35 Å², the complexes with CID 13886678 and CID 5280445 demonstrate lower and more consistent SASA values, reflecting reduced solvent exposure and suggesting that these ligands stabilize the protein’s conformation by limiting surface accessibility. Conversely, CID 41684 consistently displays the highest SASA profile among the complexes, indicating the greatest level of solvent exposure and structural variability over the simulation period. These findings underscore the stabilizing effects of CID 13886678 and CID 5280445 on the HP, with CID 13886678 showing the most consistent SASA profile, potentially indicating a more compact and less solvent-exposed conformation, which is often desirable for a potential inhibitor.

#### Principal component analysis

Principal Component Analysis (PCA) streamlines the process of pinpointing and interpreting essential coordinated movements across various protein domains (David and Jacobs 2014). This analytical approach uncovers the standout dynamic movements within simulations, spotlighting the motions that play a pivotal role in driving biological processes effectively. Known as PCA, it digs into how different elements shape the protein’s collective dance, streamlining these complex shifts to reveal their ties to stability and functionality. The visuals it produces scatter plots brimming with data points offer a window into the protein’s diverse shapes, each dot capturing a distinct structural snapshot. The simulation’s ability to reflect conformational changes over time is demonstrated by the distribution of purple and yellow dots. Time in the simulation is represented by the color progression, which goes from yellow to blue and then to purple. Here, yellow corresponds to the starting point, blue marks the midpoint, and purple indicates the concluding stage. Furthermore, the analysis indicated that the initial three principal components (PCs) represent a substantial fraction of the protein backbone dynamics evident in the MD trajectories, as illustrated in **Supplement Fig 9 and 10**. The graphic emphasizes the different properties of the complexes generated by the HP with CID5280445 and CID13886678, as shown by the PCA analysis.

The PC1 value represents the proportion of overall motion captured by the first principal component, reflecting the extent of structural fluctuations during the simulation. Among the complexes, CID 13886678 demonstrated the highest PC1 value at 55.71%; this suggests that this specific complex had the most significant structural alterations. In comparison, CID 5280445 had the lowest PC1 value at 47.47%, pointing to a relatively stable structure with minimal changes in its conformation. The control compound, CID 41684, showed a PC1 value of 56.15%, reflecting a slightly higher degree of flexibility compared to CID 13886678 and positioning it as the most dynamic among the complexes. Interestingly, the apoprotein also registered a PC1 value of 47.47%, suggesting that its unbound state displayed a similar level of movement to that of the CID 5280445 complex. The PCA plot’s clustering pattern indicates that CID 5280445 retains a more stable conformation, showing minimal deviations during the simulation. This level of stability is typically advantageous for an effective inhibitor, as significant conformational changes can signal weaker binding reliability. On the other hand, the elevated PC1 value for CID 13886678 points to increased structural variability, which might suggest greater flexibility but could also compromise the stability of the complex during formation.

#### Free Energy Landscape Analysis

To calculate the binding free energy of a protein-ligand complex, a widely used computational method known as MM/GBSA (Molecular Mechanics/Generalized Born Surface Area) was employed. This approach was applied to evaluate the binding interactions of two lead compounds with the HP, based on MD simulation trajectories. **Fig 9** illustrates that the ΔG value is determined by the combined contributions of two protein-ligand complexes and one control-protein interaction energy, including van der Waals energy (ΔEvdW), electrostatic energy (ΔEele), and EPB energy, which represents the electrostatic component of the solvation free energy calculated using the Poisson-Boltzmann method.

**Fig 9:**
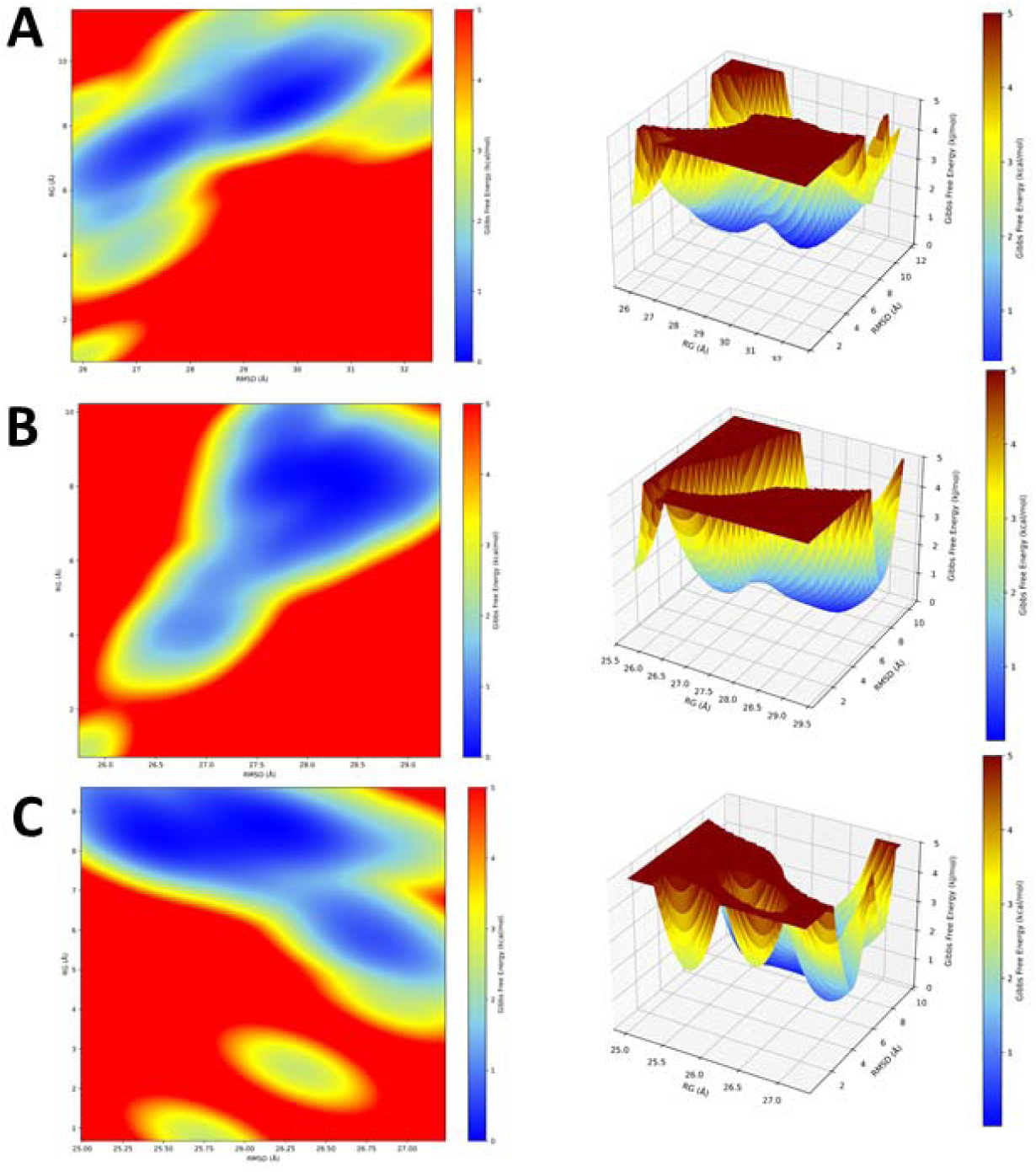
Free Energy Landscape (FEL) plots for TYC33605.1 complexes with Nitazoxanide (control), Luteolin (CID 5280445), and Sativanone (CID 13886678). Fig A (control), B (Luteolin), and C (Sativanone) show the energy landscapes, with color bars representing the corresponding free energy values. The FEL plots display the potential energy surfaces for TYC33605.1 complexes with different ligands: Nitazoxanide (A), Luteolin (B), and Sativanone (C). The color gradient represents the free energy in kcal/mol, ranging from high (red) to low (blue) energy states. Each plot corresponds to the energy distribution of the system based on the interactions between the ligand and the protein complex, highlighting the relative stability of different conformations.

The Gibbs free energy of a protein-ligand complex may be calculated by breaking down the total free energy into its component parts, which are determined by molecular mechanics and solvation energy. Given lowered Gibbs free energy (GFE), these sites are proposed to reflect thermodynamically beneficial states. Conversely, elevated RMSD values signify structural irregularities, frequently associated with decreased stability in topologies. Higher GFE in these locations indicates fewer thermodynamically favorable states and heightened energy barriers (Maisuradze et al. 2010). The troughs shown in the FEL figure mark energy minima, therefore indicating the configurations with the most stability.

#### Probability density function

The dynamic behaviors of the complexes are better understood by analyzing the Probability Density Function (PDF), which reveals patterns, trends, and structural information (Sporns et al. 2000). The PDF provides useful information on the probability of specific structural combinations happening when a bimolecular system’s structural state distribution is examined. The PDF shows the chances of seeing different sets of root-mean-square deviation (RMSD) and radius of gyration (RG) numbers. These are important measurements for checking how compact and variable a system is. There is a color bar that shows how strong these odds are. Denser bars show more common RG-RMSD pairs. Smooth distributions with distinct high-density peaks that effectively cover the required RG and RMSD ranges are characteristics of a well-defined PDF. In the analysis of the Nitazoxanide, Luteolin, and Sativanone complexes, significant PDF values were recorded within certain ranges: 32.5-11.8 Å, 29.4-10.8 Å, and 27.3-9.8 Å in the RG-RMSD space (**Fig 10**). This suggests that specific combinations of structural deviation and compactness occurred with greater frequency. Regions with higher probability densities indicate that certain conformational states are more stable or persist longer within the system. Conversely, regions with lower probability densities represent rare or transient conformational states, suggesting that the complexes are less likely to adopt these structural forms. The PDF’s contour lines make the relationship between RG and RMSD clearer. When the contour lines are closer together, it means that the link between the two qualities is stronger or that changes in one property are often accompanied by changes in the other. In contrast, if the contour lines are quite far apart, it might mean that the link between RG and RMSD is not as straightforward or that the transitions are more noticeable. This alteration in contour density emphasizes the relationship between the complexes’ temporal rigidity and structural deviation, shedding light on their structural stability and flexibility to a greater extent.

**Fig 10:**
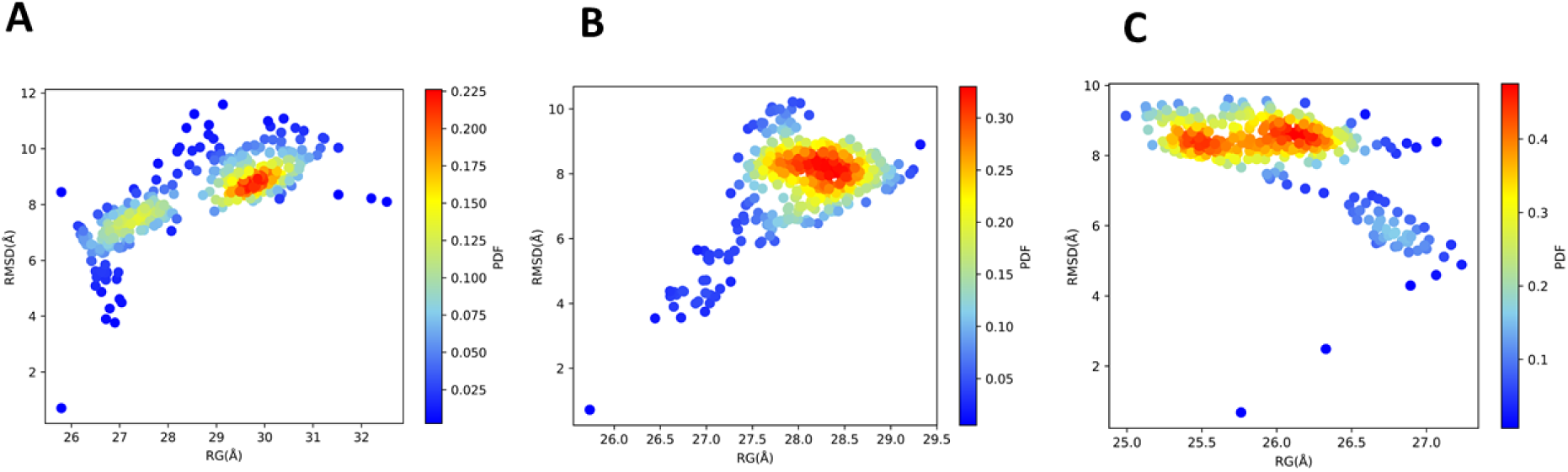
Probability density function (PDF) plots of Radius of Gyration (RG) and Root Mean Square Deviation (RMSD) distributions for TYC33605.1 complexes with Nitazoxanide (control), Luteolin (CID 5280445), and Sativanone (CID 13886678). Fig A (TYC33605.1-Nitazoxanide control), Fig B (TYC33605.1-Luteolin, CID 5280445), and Fig C (TYC33605.1-Sativanone, CID 13886678) display PDF plots comparing Radius of Gyration (RG) and Root Mean Square Deviation (RMSD) distributions, with RMSD categorized by Domain A (RMS(DA)), all-atom (RMS(AL)), and binding site (RMS(ROA)). Numerical ranges (e.g., 0.025–0.225 Å for RMSD precision; 25.0–29.5 Å for RG) under sections 1.2–5.6 reflect parameter scales across subplots, while RC(A) denotes the Reference Complex (control, Nitazoxanide).

## Conclusion

This research clarifies the structural and functional significance of the hypothetical protein TYC33605.1 in the c-di-GMP signaling-mediated biofilm formation of *Vibrio cholerae*, identified as a DGC family protein. Through integrated *in silico* approaches, including homology modeling, molecular docking, and MD simulations, Luteolin and Sativanone were identified as potent inhibitors with robust binding affinities and stable interactions with the protein’s active site. Their ability to maintain structural compactness (Rg: 25.97–27.91 Å) and reduce solvent accessibility (SASA: 25,234–25,411 Å²) underscores their potential to disrupt the c-di-GMP-mediated biofilm pathways. ADME/Tox evaluations further validated their drug-like properties and safety profiles. The findings highlight the utility of computational strategies in accelerating antimicrobial discovery, particularly against recalcitrant biofilm-associated infections. Future work should prioritize *in vitro* and *in vivo* validation of these compounds, alongside exploration of synergistic effects with existing antibiotics. This research not only advances therapeutic options for cholera but also provides a blueprint through targeting a promising HP in antibiotic resistance mechanisms.

## Acknowledgements

We are grateful to the Department of Microbiology, University of Rajshahi, for molecular dynamics support. And the Department of Biotechnology and Genetic Engineering, Gopalganj Science and Technology University, for their logistic support throughout the research.

## Author Contributions

MNHJ, MKEH, and MEH conceptualized and designed the study and conducted most of the experiments, while MSH and SS were responsible for library preparation and molecular docking. MMHS, MFH, and AKD carried out the molecular dynamics (MD) simulations. Data analysis was performed by MNHJ, MKEH, and MEH. MEH contributed to specific experiments and provided valuable guidance for the study. MNHJ, MKEH, MSH, SS, AS, and MEH drafted the manuscript, which all authors reviewed and approved for publication.

## Abbreviation

HP: hypothetical protein,
DGC: Diguanylate cyclase,
c-di-GMP: cyclic-di-GMP,
QS: quorum sensing,
MD: Molecular Dynamics,
DFT: Density Functional Theory,
PD: Pharmacodynamics,
PK: Pharmacokinetics,
PCA: Principal Component Analysis,
PDF: Probability Density Function,
FEL: Free Energy Landscape.

