## Supplement Fig 1 for "Computational Structure Modeling, Functional Characterization, and Identification of Potential Inhibitors for the cyclic-di-GMP Mediated Biofilm Forming Membrane Protein in *Vibrio cholerae*"

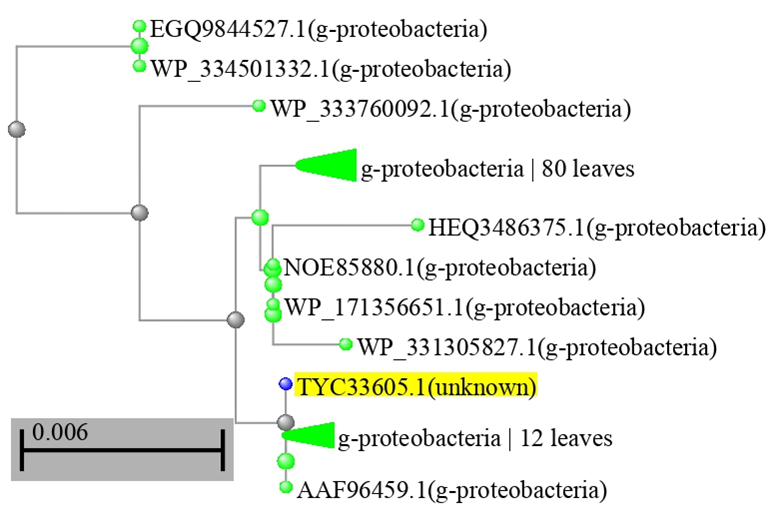
**Supplement Fig 1: Phylogenetic structures based on the genetic relatedness of the hypothetical protein**. The figure depicts the alignment or structural comparison of two sequences: TYC33605.1 (unidentified origin) and WP_053032761.1 (NCBI protein accession). Conserved regions are highlighted in blue, divergent regions in orange, and annotations reflect standard database conventions. WP_053032761.1 represents a characterized protein entry in the NCBI non-redundant database, while TYC33605.1 lacks explicit functional annotation.
