## Supplement Fig 2 for "Computational Structure Modeling, Functional Characterization, and Identification of Potential Inhibitors for the cyclic-di-GMP Mediated Biofilm Forming Membrane Protein in *Vibrio cholerae*"

**Supplement Fig 2: Amino acid composition and physicochemical properties of TYC33605.1. (A) Bar plot of amino acid frequencies. (B) Key parameters: pI, instability index, and hydropathicity.**

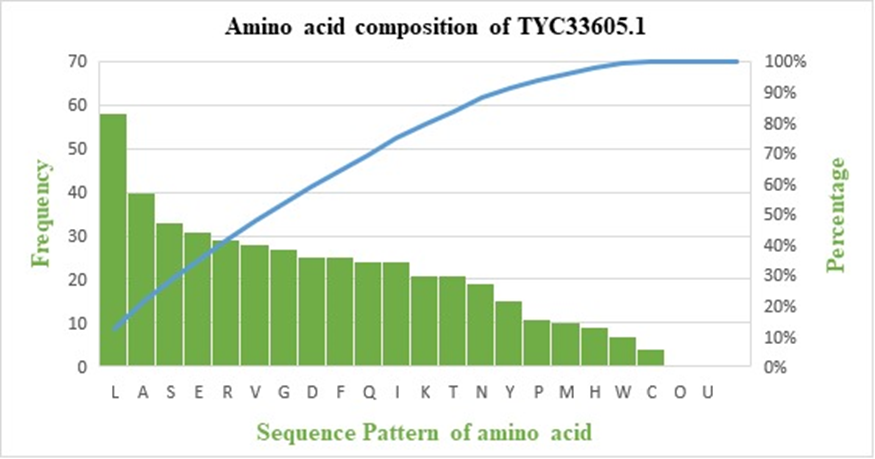
