## Supplement Fig 3 for "Computational Structure Modeling, Functional Characterization, and Identification of Potential Inhibitors for the cyclic-di-GMP Mediated Biofilm Forming Membrane Protein in *Vibrio cholerae*"

**Supplement Fig 3: Coiled-coil probability plot of TYC33605.1 generated using the COILS server**. Peaks indicate regions with high coiled-coil propensity, analyzed using window sizes 14 (red), 21 (green), and 28 (blue)

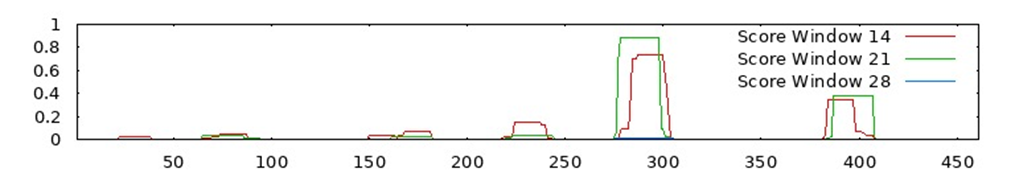
