## Supplement Fig 4 for "Computational Structure Modeling, Functional Characterization, and Identification of Potential Inhibitors for the cyclic-di-GMP Mediated Biofilm Forming Membrane Protein in *Vibrio cholerae*"

**Supplement Fig 4: Secondary structure prediction of TYC33605.1 using PSIPRED and SOPMA**. Predicted α-helices (53.36%), β-strands (15.84%), and random coils (30.80%) are shown.

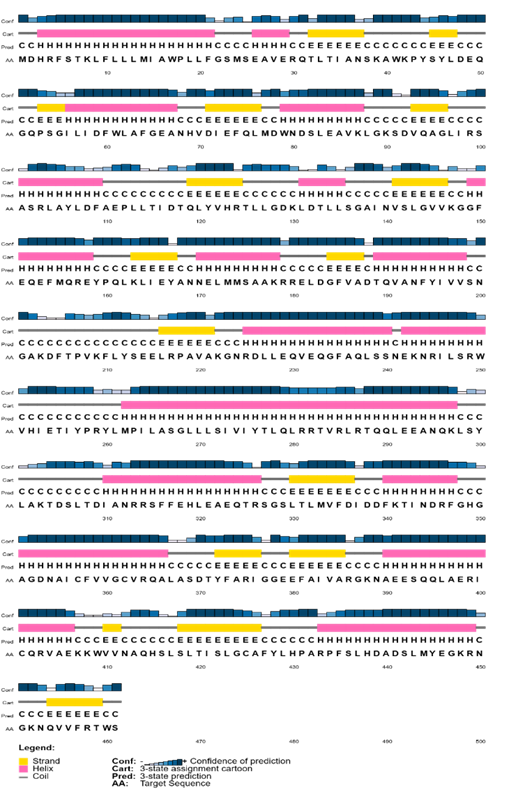
