## Supplement Fig 5 for "Computational Structure Modeling, Functional Characterization, and Identification of Potential Inhibitors for the cyclic-di-GMP Mediated Biofilm Forming Membrane Protein in *Vibrio cholerae*"

**Supplement Fig 5: Ramachandran plot of the refined TYC33605.1 model validated using MolProbity**. 91.1% of residues lie in favored regions, confirming stereochemical quality.

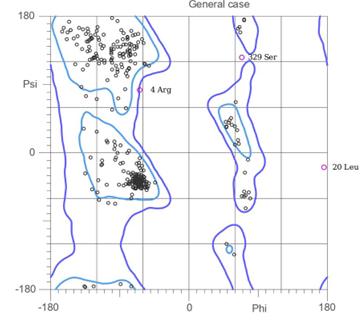
