## Supplement Fig 6 for "Computational Structure Modeling, Functional Characterization, and Identification of Potential Inhibitors for the cyclic-di-GMP Mediated Biofilm Forming Membrane Protein in *Vibrio cholerae*"

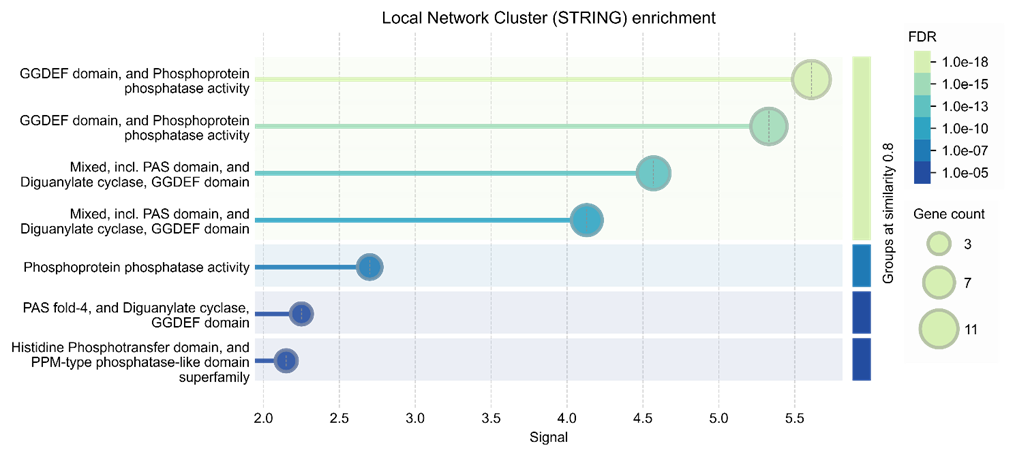
**Supplement Fig 6. Functional enrichment analysis of protein-protein interaction network conducted using the STRING database**. The bubble chart represents significantly enriched Gene Ontology (GO) biological processes associated with the protein set. The x-axis denotes the enrichment significance (−log10 FDR), while bubble size reflects the number of genes involved in each GO term. The color gradient indicates the false discovery rate (FDR), with darker shades representing higher statistical significance. The analysis reveals key biological processes such as [insert top GO terms if desired], suggesting functional similarity and potential interactions among the analyzed proteins.
