## Supplement Fig 7 for "Computational Structure Modeling, Functional Characterization, and Identification of Potential Inhibitors for the cyclic-di-GMP Mediated Biofilm Forming Membrane Protein in *Vibrio cholerae*"

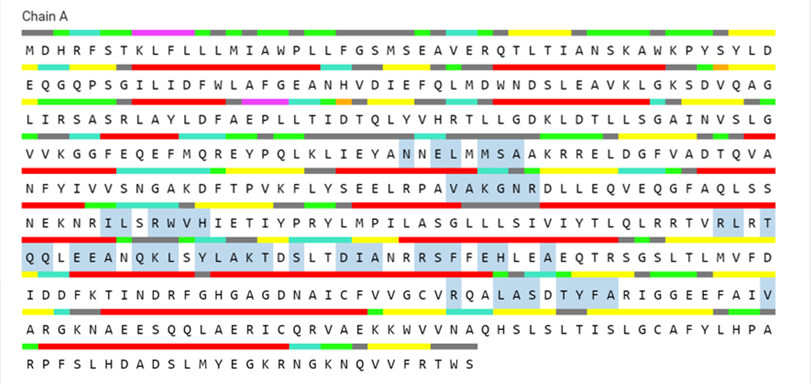
**Supplementary Fig 7: Active site residues of TYC33605.1 predicted by CASTp**. Key residues (Asn169, Glu293, Arg364, Phe373) form the ligand-binding pocket (blue surface). Annotations highlight potential drug-target interactions.
