## Supplement Fig 8 for "Computational Structure Modeling, Functional Characterization, and Identification of Potential Inhibitors for the cyclic-di-GMP Mediated Biofilm Forming Membrane Protein in *Vibrio cholerae*"

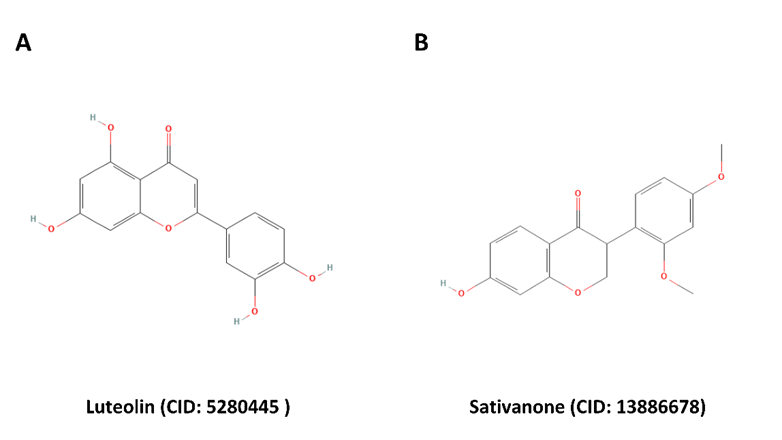
**Supplementary Fig 8: Molecular structures of Luteolin (CID: 5280445) and Sativanone (CID: 13886678).** The diagram highlights key atomic components and functional groups: A and B denote distinct structural regions (e.g., aromatic rings or side chains), H (hydrogen) and O (oxygen) atoms mark hydroxyl (-OH) and carbonyl (C=O) groups critical to their bioactive properties. Luteolin, a flavonoid, features multiple hydroxyl groups on its benzopyran backbone, while Sativanone, an isoflavanone, includes a prenylated moiety (B) and oxygen-rich functional groups, reflecting their roles in antioxidant and antimicrobial interactions
