## Supplement Fig 9 for "Computational Structure Modeling, Functional Characterization, and Identification of Potential Inhibitors for the cyclic-di-GMP Mediated Biofilm Forming Membrane Protein in *Vibrio cholerae*"

**Supplement Fig 9: PCA plot of TYC33605.1 complex with Luteolin (CID 5280445).** The scatter plots show the distribution along the first three principal components (PC1, PC2, and PC3), with color indicating variance explained by each component. The cumulative variance plot is shown on the right.

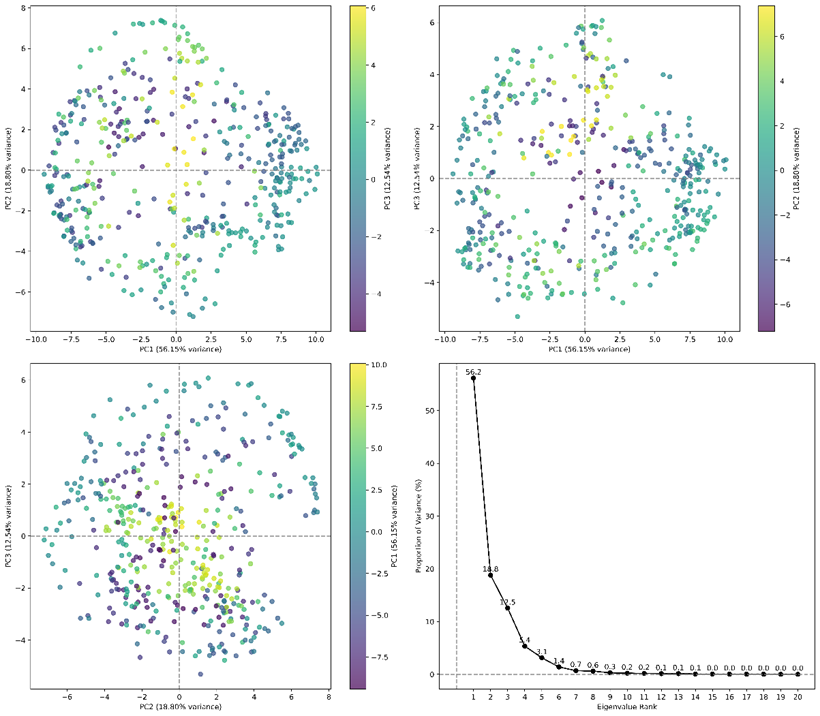
