## Supplement Fig 10 for "Computational Structure Modeling, Functional Characterization, and Identification of Potential Inhibitors for the cyclic-di-GMP Mediated Biofilm Forming Membrane Protein in *Vibrio cholerae*"

**Supplement Fig 10: PCA plot of TYC33605.1 complex with Sativanone (CID 13886678).** The scatter plots show the distribution along the first three principal components (PC1, PC2, and PC3), with color indicating variance explained by each component. The cumulative variance plot is shown on the right.

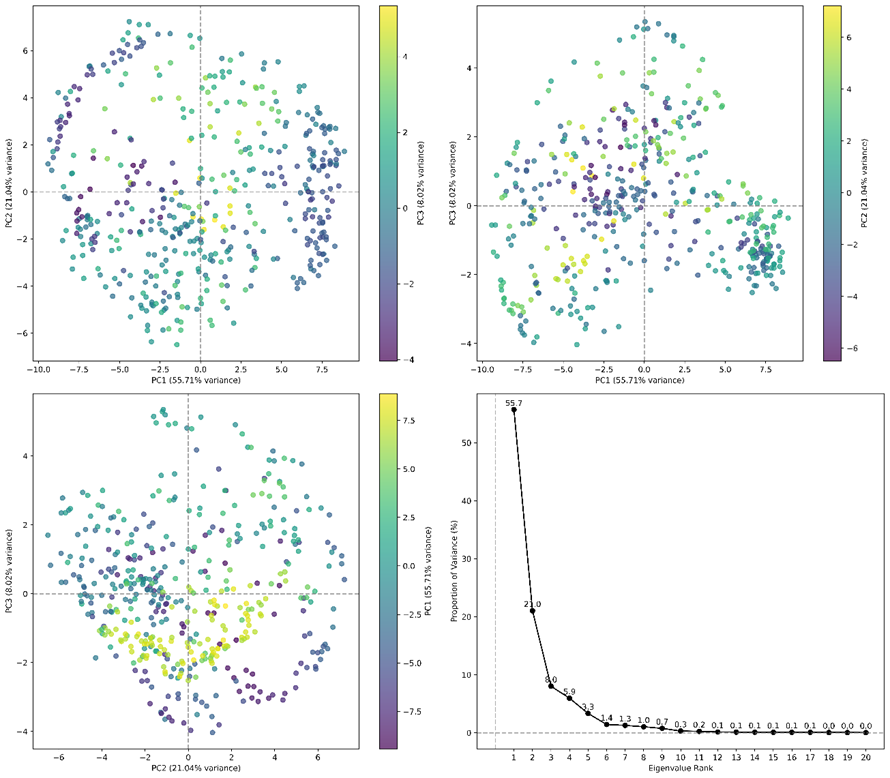
