## Supplement Table 1 for "Computational Structure Modeling, Functional Characterization, and Identification of Potential Inhibitors for the cyclic-di-GMP Mediated Biofilm Forming Membrane Protein in *Vibrio cholerae*"

**Supplement Table 1: Bioinformatics tools and servers used for functional annotation, structural modeling, and molecular dynamics simulations of the hypothetical protein**

| **Serial** | **Tools** | **Server URL** | **Application** |
| --- | --- | --- | --- |
| 1 | NCBI protein Database | <https://www.ncbi.nlm.nih.gov/> | Retrieve protein sequence |
| 2 | Clustal Omega | <https://www.ebi.ac.uk/Tools/msa/clustalo/> | Sequence Alignment and Tree Construction |
| 3 | MEGA X [21] | <https://www.megasoftware.net/> |  |
| 4 | iTOL server [22] | <https://itol.embl.de/> |  |
| 5 | ProtParam ExPASy | <https://web.expasy.org/protparam/> | Physicochemical characterization |
| 6 | CELLO server | <https://cello.life.nctu.edu.tw/> | Subcellular localization prediction |
| 7 | PSORTb | <https://psort.org/psortb/> |  |
| 8 | PSLpred | <https://crdd.osdd.net/raghava/pslpred/> |  |
| 9 | SOSUIGramN | <https://harrier.nagahama-i-bio.ac.jp/sosui/sosuigramn/sosuigramn_submit.html> |  |
| 10 | TMHMM 2.0 | <https://services.healthtech.dtu.dk/services/TMHMM-2.0/> | Topology prediction |
| 11 | HMMTOP | <https://hmmtop.pbrg.hu/> |  |
| 12 | CCTOP | <https://cctop.ttk.hu/> |  |
| 13 | Genome Net server | <https://www.genome.jp/tools/motif/> | Motif discovery |
| 14 | Pfam | <https://pfam.xfam.org> | Protein Domain Discovery |
| 15 | NCBI’s conserved domain database (CDD) | <https://www.ncbi.nlm.nih.gov/Structure/cdd/wrpsb.cgi> | Conserved domain analysis |
| 16 | InterProScan server | <https://www.ebi.ac.uk/interpro/search/sequence> | Functional classification |
| 17 | PFP-FunD SeqE server | <http://www.csbio.sjtu.edu.cn/bioinf/PFP-FunDSeqE> | Fold recognition |
| 18 | COILS server | <https://npsa-pbil.ibcp.fr/cgi-bin/npsa_automat.pl?page=/NPSA/npsa_lupas.html> | Coiled-coil motif identification |
| 19 | DISULFIND server [23] | <http://disulfind.dsi.unifi.it/> | Stability of folding process analysis |
| 20 | STRING 12.0 | <https://string-db.org> | Protein Interactions Analysis |
| 21 | PSIPRED | <http://bioinf.cs.ucl.ac.uk/psipred/> | Secondary structure prediction |
| 22 | SOPMA | <https://npsa-pbil.ibcp.fr/cgi-bin/npsa_automat.pl?page=/NPSA/npsa_sopma.html> |  |
| 23 | I-TASSER | <https://zhanggroup.org/I-TASSER> | Tertiary structure prediction and Visualization |
| 24 | PyMOL | <https://www.pymol.org/> |  |
| 25 | ModRefiner | <http://zhanglab.ccmb.med.umich.edu/ModRefiner/> | Three-Dimensional Structure Refinement |
| 26 | MolProbity | <http://molprobity.biochem.duke.edu/> | Model quality assessment |
| 27 | Verify3D | <http://nihserver.mbi.ucla.edu/Verify_3D/> |  |
| 28 | ERRAT | <https://servicesn.mbi.ucla.edu/ERRAT/> |  |
| 29 | CASTp | <http://sts.bioengr.uic.edu/castp/> | Active site prediction |
