## Supplement Table 2 for "Computational Structure Modeling, Functional Characterization, and Identification of Potential Inhibitors for the cyclic-di-GMP Mediated Biofilm Forming Membrane Protein in *Vibrio cholerae*"

**Supplement Table 2: Homologous proteins to TYC33605.1 identified in the UniProt database, highlighting conserved functional domains**

| Protein ID | Organism | Protein name | Identity (%) | Score | e-value |
| --- | --- | --- | --- | --- | --- |
| Q0T466.1 | Shigella flexneri | Diguanylate cyclase (DGC) | 36.47% | 48.9 | 5e-05 |
| Q83KV7.1 | Shigella flexneri | Diguanylate cyclase (DGC) | 36.47% | 48.9 | 5e-05 |
| P54595.1 | Bacillus subtilis | Uncharacterized protein YhcK | 36.09% | 95.1 | 3e-20 |
| Q9KKZ4.1 | Vibrio cholerae | Diguanylate cyclase (DGC) | 35.56% | 72.4 | 1e-12 |
| P75908.1 | Escherichia coli | Probable Diguanylate cyclase (DGC) | 35.54% | 97.4 | 1e-20 |
| P64826.1 | Mycobacterium tuberculosis | Uncharacterized protein Mb1389c | 35.29% | 80.9 | 6e-15 |
| P76245.2 | Escherichia coli | Diguanylate cyclase (DGC) | 34.71% | 70.5 | 5e-12 |
| A0A0H2ZJS2.1 | Pseudomonas aeruginosa | Diguanylate cyclase (DGC) | 34.54% | 95.1 | 1e-19 |
| Q9HT84.1 | Pseudomonas aeruginosa | Diguanylate cyclase (DGC) | 34.54% | 94.4 | 2e-19 |
| Q55434.1 | Synechocystis sp. | Phytochrome-like protein cph2 | 34.29% | 81.6 | 5e-15 |
