## Supplement Table 3 for "Computational Structure Modeling, Functional Characterization, and Identification of Potential Inhibitors for the cyclic-di-GMP Mediated Biofilm Forming Membrane Protein in *Vibrio cholerae*"

**Supplement Table 3: Subcellular localization predictions for TYC33605.1 using multiple tools**

| **No.** | **Analysis** | **Result** |
| --- | --- | --- |
| 01 | CELLO 2.5 | Inner Membrane, Cytoplasmic |
| 02 | PSORTb | Periplasmic Protein |
| 03 | SOSUIGramN | IM (inner membrane) |
| 04 | PSLpred | Cytoplasmic Membrane |
| 05 | TMHMM 2.0 | One transmembrane helices present |
| 06 | HMMTOP | Two transmembrane helices present |
| 07 | CCTOP | One transmembrane helices present |
