## Supplement Table 4 for "Computational Structure Modeling, Functional Characterization, and Identification of Potential Inhibitors for the cyclic-di-GMP Mediated Biofilm Forming Membrane Protein in *Vibrio cholerae*"

**Supplement Table 4: Domain architecture of TYC33605.1 identified via CDD-BLAST and Pfam**

| Name | Accession | Description | Interval (Amino acid) | E-value |
| --- | --- | --- | --- | --- |
| PBP2_HisK_like_1 | cd13706 | Putative sensor domain similar to HisK: includes periplasmic sensor domain of the histidine kinase receptors (HisK) which are elements of the two-component signal transduction systems | 31-251 | 4.81e-99 |
| GGDEF | COG2199 | GGDEF domain, diguanylate cyclase (c-di-GMP synthetase), is a bacterial second messenger that regulates cell surface-associated traits such as biofilm formation, cellulose production, and motility. | 191-457 | 8.32e-53 |
| GGDEF | cd01949 | Diguanylate-cyclase (DGC) or GGDEF domain. | 304-456 | 1.31e-49 |
| SBP_bac_3 | pfam00497 | Bacterial extracellular solute-binding proteins: a sensor domain found in solute-binding protein family from Gram-positive bacteria, Gram-negative bacteria, and archaea | 33-250 | 2.12e-33 |
