## Supplement Table 5 for "Computational Structure Modeling, Functional Characterization, and Identification of Potential Inhibitors for the cyclic-di-GMP Mediated Biofilm Forming Membrane Protein in *Vibrio cholerae*"

**Supplement Table 5: Motifs identified in TYC33605.1 using Pfam and Genome Net servers.**

| Pfam | Position (Independent E-value) | Description |
| --- | --- | --- |
| GGDEF | 302-454 (7.8 x 10^-34^) | Diguanylate cyclase, GGDEF domain (PF00990) |
| SBP_bac_3 | 33-251 (4.1 x 10^-19^) | Bacterial extracellular solute-binding proteins (PF00497) |
