## Supplement Table 6 for "Computational Structure Modeling, Functional Characterization, and Identification of Potential Inhibitors for the cyclic-di-GMP Mediated Biofilm Forming Membrane Protein in *Vibrio cholerae*"

### Supplementary Table 6: Collected 1092 compounds from 17 medicinal plants.

| Serial no | Plants Name | Compound name | Pubchem id | Ref |
| --- | --- | --- | --- | --- |
| 1 | Acacia nilotica L | 4-methylbenzenethiol | 7811 | [1] |
|  |  | Pyrogallol | 1057 |  |
|  |  | 1,8,11-Heptadecatriene, (Z,Z)- | 5352709 |  |
|  |  | 4-O methylmannose | 345716 |  |
|  |  | Hexadecanoic acid, methyl ester | 8181 |  |
|  |  | 14,17-Octadecadienoic acid, methyl ester | 5365751 |  |
|  |  | 9,12-Octadecadienoic acid (Z,Z)- | 5280450 |  |
|  |  | Methyl oleate | 5364509 |  |
|  |  | Methyl linoleate | 5284421 |  |
|  |  | Methyl 9-cis,11-trans-octadecadienoate | 11748436 |  |
|  |  | Methyl stearate | 8201 |  |
|  |  | 15-Hydroxypentadecanoic acid | 78360 |  |
|  |  | Oxiranyl methyl ester 9-octadecenoic acid | 5354568 |  |
|  |  | 9-Octadece mide | 1930 |  |
|  |  | Phthalic acid, bis(2-ethylhexyl) ester | 8343 |  |
|  |  | Ergosta-5,22-dien-3-ol, (3.beta.,22E)- | 6432458 |  |
|  |  | Ergost-5-en-3-ol | 18660356 |  |
|  |  | Decane, 3,7-dimethyl- | 28468 | [2] |
|  |  | Dihydrocitronellol | 7792 |  |
|  |  | Pelargo ldehyde | 31289 |  |
|  |  | Undecane | 14257 |  |
|  |  | No ne, 5-(2-methylpropyl)- | 545936 |  |
|  |  | 1-Chlorohexadecane | 20993 |  |
|  |  | Hexadecane | 11006 |  |
|  |  | 1,3,4-Eugenol | 3314 |  |
|  |  | Pentadecane | 12391 |  |
|  |  | Phosphoric acid, bis(trimethylsilyl)monomethyl ester | 617748 |  |
|  |  | Heptadecane | 12398 |  |
|  |  | 3',5'-Dimethoxyacetophenone | 95997 |  |
|  |  | Fumaric acid, ethyl 2-methylallyl ester | 5461492 |  |
|  |  | Phthalic acid | 1017 |  |
|  |  | Megastigmatrienone | 5375190 |  |
|  |  | 4-(1,5-Dihydroxy-2,6,6-trimethylcyclohex-2-enyl)but-3-en-2-one | 5363693 |  |
|  |  | 3-Oxo-.alpha.-ionol | 5370052 |  |
|  |  | (3-Oxo-2-pent-2-enylcyclopentyl)acetic acid | 5367720 |  |
|  |  | Myristic acid | 11005 |  |
|  |  | 2-Methylresorcinol, acetate | 23469416 |  |
|  |  | Neophytadiene | 10446 |  |
|  |  | Tetracosane | 12592 |  |
|  |  | 1,54-Dibromotetrapentacontane | 545963 |  |
|  |  | Isopropyl Palmitate | 8907 |  |
|  |  | 1,11-Hexadecadiyne | 3085574 |  |
|  |  | Linolenic acid, methyl ester | 5319706 |  |
|  |  | Cedrane-8,13-diol | 188457 |  |
|  |  | Tetracosane | 12592 |  |
|  |  | Linolenic acid | 5280934 |  |
|  |  | Stearic acid | 5281 |  |
|  |  | Stearic acid ethyl ester | 8122 |  |
|  |  | Dotriacontane | 11008 |  |
|  |  | Arachidonic acid | 444899 |  |
|  |  | Dipalmitin | 71308681 |  |
|  |  | Octadecane, 3-ethyl-5-(2-ethylbutyl)- | 292285 |  |
|  |  | Tetracosane | 12592 |  |
|  |  | 1,3,5-Trisilacyclohexane | 6329365 |  |
|  |  | Palmitoyl chloride | 8206 |  |
|  |  | Phenol, 4,4'-methylenebis[2,6-bis(1,1-dimethylethyl)- | 8372 |  |
|  |  | Tetrapentacontane | 521846 |  |
|  |  | Decyl sulfide | 69662 |  |
|  |  | Hexatriacontane | 12412 |  |
|  |  | Oxirane, hexadecyl- | 23872 |  |
|  |  | Hexatriacontane | 12412 |  |
|  |  | Tetrapentacontane | 521846 |  |
|  |  | (+)-Lariciresinol | 332427 |  |
|  |  | (+)-Lariciresinol | 332427 |  |
|  |  | 1,3,4-Eugenol | 3314 |  |
|  |  | Lariciresinol | 332427 |  |
|  |  | Phthalic acid | 1017 |  |
|  |  | N-2,4-Dnp-L-arginine | 7083742 | [3] |
|  |  | Pyridine-3-carboxamide | 936 |  |
|  |  | Benzaldehyde, 4-methyl- | 7725 |  |
|  |  | Tridecane, 1-bromo- | 13000 |  |
|  |  | Phenylacetic acid, 2-adamantyl ester | 584925 |  |
|  |  | Benzene, 1-(3-trifluoromethylphenyliminomethyl)-4-(thietan-3-yloxy)- | 552166 |  |
|  |  | Carbonic acid, ethyl phenyl ester | 138067 |  |
|  |  | Benzene, 1,3-bis(1,1-dimethylethyl)- | 136810 |  |
|  |  | 4-Trifluoroacetoxytridecane | 543275 |  |
|  |  | 2-Trifluoroacetoxypentadecane | 534405 |  |
|  |  | 1-Octanol, 2-butyl- | 19800 |  |
|  |  | 1-Dodecanol | 8193 |  |
|  |  | Phenol, 2,5-bis(1,1-dimethylethyl)- | 79983 |  |
|  |  | Silane, (dodecyloxy)trimethyl- | 22587 |  |
|  |  | Dodecyl acrylate | 75084 |  |
|  |  | Propanoic acid, decyl ester | 229385 |  |
|  |  | Dodecanoic acid, 2,3-bis(acetyloxy)propyl ester | 169212 |  |
|  |  | 4-CyclopropylcarbonyloxytetraDecane | 549497 |  |
|  |  | Pyridine-3-carboxamide | 936 |  |
|  |  | Tetracontane, 3,5,24-trimethyl- | 41344 |  |
|  |  | Decane, 1-(ethenyloxy)- | 12999 |  |
|  |  | Cis-9-Hexadecenoic acid | 445638 |  |
|  |  | Pentadecanoic acid, 14-methyl-, methyl ester | 21205 |  |
|  |  | Dibutyl phthalate | 3026 |  |
|  |  | Octadeca l | 12533 |  |
|  |  | Propanoic acid, 3-mercapto-, dodecyl ester | 80796 |  |
|  |  | Propionic acid, 3-mercapto-, 2,2,4,4-tetramethylpentyl ester | 545899 |  |
|  |  | Tridecane, 1-bromo- | 13000 |  |
|  |  | Dodecyl acrylate | 75084 |  |
|  |  | Tetracontane, 3,5,24-trimethyl- | 41344 |  |
|  |  | Cis-9-Hexadecenoic acid | 445638 |  |
|  |  | 2-Pentadecanone, 6,10,14-trimethyl- | 10408 |  |
|  |  | Hexadecanoic acid, methyl ester | 8181 |  |
|  |  | Dibutyl phthalate | 3026 |  |
|  |  | Cis,cis,cis-7,10,13-Hexadecatrie l | 5367366 |  |
|  |  | Hexadecane | 11006 |  |
| 2 | Aegle marmelos | Lupeol | 259846 | [4] |
|  |  | acetamide | 178 |  |
|  |  | benzoic acid | 243 |  |
|  |  | pyranocoumarin | 10606 |  |
|  |  | scopoletin | 5280460 |  |
|  |  | marmesin | 334704 |  |
|  |  | psoralen | 6199 |  |
|  |  | skimmianine | 6760 |  |
|  |  | eugenol | 3314 |  |
| 3 | Andrographis paniculata | 1,4 Dichlorobenzene | 4685 | [5] |
|  |  | 1-Dodecene | 8183 |  |
|  |  | 2-Tetradecene | 33650 |  |
|  |  | Pentadecane | 12391 |  |
|  |  | Eicosane | 8222 |  |
|  |  | Benzofuranone | 68382 |  |
|  |  | 1-Hexadecene | 12395 |  |
|  |  | Diethyl phthalate | 6781 |  |
|  |  | Heptadecane | 12398 |  |
|  |  | Pentadecane | 12391 |  |
|  |  | Heptacosane | 11636 |  |
|  |  | 5-Octadecene | 5364598 |  |
|  |  | Octadecane | 11635 |  |
|  |  | Neophytadiene | 10446 |  |
|  |  | Pentadecanoic acid, methyl ester | 23518 |  |
|  |  | Hexadecanoic acid, ethyl ester | 12366 |  |
|  |  | Tricosane | 12534 |  |
|  |  | Eicosane | 8222 |  |
|  |  | Tetracosane | 12592 |  |
|  |  | Pentacosane | 12406 |  |
|  |  | Di-n-octyl phthalate | 8346 |  |
|  |  | Hexacosane | 12407 |  |
|  |  | Heneicosane | 12403 |  |
|  |  | Allyl acetate | 11584 | [6] |
|  |  | 2-Monolinolenin, 2TMS derivative | 5362857 |  |
|  |  | p-Cresyl glycidyl ether | 16606 |  |
|  |  | 2,6-Dimethoxy-4-(2-propenyl)-phenol | 226486 |  |
|  |  | Oleic acid | 445639 |  |
|  |  | Dibutyl phthalate | 3026 |  |
|  |  | Phytol | 5280435 |  |
|  |  | 4-Ethylbenzoic acid, 4-hexadecyl ester | 530442 |  |
|  |  | Hexadecanoic acid, methyl ester | 8181 |  |
|  |  | 9-Octadecenoic acid (Z)-, methyl ester | 5364509 |  |
|  |  | 2,5-Octadecadiynoic acid, methyl ester | 42151 |  |
| 4 | Artemisia absinthium L | Alpha-Thujene | 17868 | [7] |
|  |  | Alpha-Pinene | 6654 |  |
|  |  | Camphene | 6616 |  |
|  |  | Sabinene | 18818 |  |
|  |  | Beta-Pinene | 14896 |  |
|  |  | Myrcene | 31253 |  |
|  |  | Alpha-Phellandrene | 7460 |  |
|  |  | Alpha-Terpinene | 7462 |  |
|  |  | p-Cymene | 7463 |  |
|  |  | Eucalyptol | 2758 |  |
|  |  | Limonene | 22311 |  |
|  |  | cis-Arbusculone | 21630868 |  |
|  |  | Artemisia ketone | 68346 |  |
|  |  | Gamma-Terpinene | 7461 |  |
|  |  | 4-Thujanol | 62367 |  |
|  |  | 2-No none | 13187 |  |
|  |  | Terpinolene | 11463 |  |
|  |  | Li lool | 6549 |  |
|  |  | trans-Sabinene hydrate | 6326181 |  |
|  |  | Camphor | 2537 |  |
|  |  | Camphene hydrate | 101680 |  |
|  |  | Borneol | 64685 |  |
|  |  | Terpinen-4-ol | 11230 |  |
|  |  | Dill ether | 126537 |  |
|  |  | Alpha-Terpineol | 17100 |  |
|  |  | Dihydrocarvone | 24473 |  |
|  |  | Carvone | 7439 |  |
|  |  | Bornyl acetate | 93009 |  |
|  |  | Ethyl hydrocin mate | 16237 |  |
|  |  | (Z)-Ethyl cin mate | 5284656 |  |
|  |  | (E)-Methyl cin mate | 637520 |  |
|  |  | Biphenyl | 7095 |  |
|  |  | (E)-Beta-Caryophyllene | 5281515 |  |
|  |  | (E)-Beta-Farnesene | 5281517 |  |
|  |  | (E)-Ethyl cin mate | 637758 |  |
|  |  | Beta-Selinene | 442393 |  |
|  |  | Dava ether | 5370105 |  |
|  |  | (E)-Nerolidol | 5284507 |  |
|  |  | Spathulenol | 92231 |  |
|  |  | Beta-Eudesmol | 91457 |  |
|  |  | Davanol | 529918 |  |
|  |  | Chamazulene | 10719 |  |
|  |  | Margaspidin | 15854 | [8] |
|  |  | Stigmasterol | 5280794 |  |
|  |  | Octadecanoic acid, 2,3-dihydroxypropyl ester | 24699 |  |
|  |  | Hexadecanoic acid, 2,3-dihydroxypropyl ester | 14900 |  |
|  |  | 7-Hexadecyn-1-ol | 549047 |  |
|  |  | 2-Propenoic acid, 3-phenyl-, ethyl ester | 637758 |  |
| 5 | Chlorophytum borivilianum L. | Oleic acid | 445639 | [9] |
|  |  | Xanthotoxin | 4114 |  |
|  |  | Stearic acid | 5281 |  |
|  |  | Heptadecanoic acid | 10465 |  |
|  |  | Behenic acid | 8215 |  |
|  |  | Stigmasterol | 5280794 |  |
|  |  | 1-hexadecanol | 2682 |  |
|  |  | Sucrose | 5988 |  |
|  |  | Lauric acid | 3893 |  |
|  |  | Valine | 6287 | [10] |
|  |  | Thymine | 1135 |  |
|  |  | 4-mercaptophenol | 240147 |  |
|  |  | Hexadecanoic acid, 3-hydroxy-, methyl ester | 103553 |  |
|  |  | Hexadeca l | 984 |  |
|  |  | Octadeca l | 12533 |  |
|  |  | Pentadeca l | 17697 |  |
|  |  | Phytol | 5280435 |  |
|  |  | Dodecanoic acid | 3893 |  |
|  |  | Undecanoic acid | 8180 |  |
|  |  | 9,12-Octadecadienoic acid | 3931 |  |
|  |  | 9,12,15-octadecatrienoic acid | 860 |  |
|  |  | Gamolenic acid | 5280933 |  |
|  |  | N-Decanoic acid | 2969 |  |
|  |  | Triarachine | 522017 |  |
|  |  | Palmitoyl chloride | 8206 |  |
|  |  | Betulin | 72326 |  |
|  |  | Taraxasterol | 115250 |  |
|  |  | Levomenthol | 16666 |  |
| 6 | Cinnamomum verum L. | Alpha-Pinene | 6654 | [11] |
|  |  | Camphene | 6616 |  |
|  |  | Benzaldehyde | 240 |  |
|  |  | Beta-Pinene | 14896 |  |
|  |  | 6-Methyl-5-hepten-2-one | 9862 |  |
|  |  | p-Cymene | 7463 |  |
|  |  | D-Limonene | 440917 |  |
|  |  | Salicylaldehyde | 6998 |  |
|  |  | Phenethyl Alcohol | 6054 |  |
|  |  | Phenylpropyl Aldehyde | 7707 |  |
|  |  | Borneol | 64685 |  |
|  |  | Cin mic Alcohol | 5315892 |  |
|  |  | Alpha-Terpineol | 17100 |  |
|  |  | Cin mic Aldehyde | 637511 |  |
|  |  | o-Anisaldehyde | 8658 |  |
|  |  | Phenyl Ethyl Acetate | 20354 |  |
|  |  | Trans-2-dece l | 5283345 |  |
|  |  | Cyclosativene | 519960 |  |
|  |  | Alpha-Ylangene | 442409 |  |
|  |  | Alpha-Copaene | 19725 |  |
|  |  | Eugenol | 3314 |  |
|  |  | Beta-Elemene | 6918391 |  |
|  |  | Caryophyllene | 5281515 |  |
|  |  | Alpha-Himachalene | 11830551 |  |
|  |  | Cin myl Acetate | 5282110 |  |
|  |  | 2-Methoxycin maldehyde | 641298 |  |
|  |  | Gamma-Muurolene | 12313020 |  |
|  |  | Alpha-Muurolene | 12306047 |  |
|  |  | Alpha-Curcumene | 92139 |  |
|  |  | Beta-Bisabolene | 10104370 |  |
|  |  | Delta-Cadinene | 441005 |  |
|  |  | Alpha-Longipinene | 12311396 |  |
|  |  | Nerolidol | 5284507 |  |
|  |  | Spathulenol | 92231 |  |
|  |  | Globulol | 12304985 |  |
|  |  | Benzyl Benzoate | 2345 |  |
|  |  | Benzaldehyde | 240 | [12] |
|  |  | Benzeneacetaldehyde | 998 |  |
|  |  | Acetophenone | 7410 |  |
|  |  | Benzenepropa l | 7707 |  |
|  |  | (Z)-3-Phenylacrylaldehyde | 6428995 |  |
|  |  | Hydrocin mic acid | 107 |  |
|  |  | 3-phenyl-2-Propenoic acid | 444539 |  |
|  |  | (Z)-Cin myl acetate | 5315912 |  |
|  |  | trans-Cin mic acid | 444539 |  |
|  |  | Cis-Cin mic acid | 5372954 |  |
|  |  | 3-(2-methoxyphenyl)-2-prope l | 641298 |  |
|  |  | Tetradeca l | 31291 |  |
|  |  | Chalcone | 637760 |  |
|  |  | 1,4-diphenyl-1,4-butanedione | 136322 |  |
|  |  | 5-Phenyl-2,4-pentadienophenone | 1549519 |  |
|  |  | Beta-amyrin | 73145 |  |
|  |  | P-cymene | 7463 | [13] |
|  |  | Heptanoic acid | 8094 |  |
|  |  | Benzenepropa l | 7707 |  |
|  |  | Li lool | 6549 |  |
|  |  | Li lyl propio te | 61098 |  |
|  |  | Borneol | 64685 |  |
|  |  | Cis-cin maldehyde | 6428995 |  |
|  |  | Trans-cin maldehyde | 637511 |  |
|  |  | Eugenol | 3314 |  |
|  |  | Copaene | 12303902 |  |
|  |  | Cin myl acetate | 5282110 |  |
|  |  | Trans-caryophyllene | 5281515 |  |
|  |  | Cin mic acid | 444539 |  |
|  |  | Alpha-Humulene | 5281520 |  |
|  |  | o-Methoxycin maldehyde | 641298 |  |
|  |  | Ledene | 10910653 |  |
|  |  | Gurjunene | 15560275 |  |
|  |  | Alpha-Cadinene | 12306048 |  |
|  |  | Spathulenol | 92231 |  |
|  |  | Clovene | 10102 |  |
|  |  | Humuladienone | 101297706 |  |
|  |  | Bisabolol | 1549992 |  |
|  |  | Benzyl benzoate | 2345 |  |
|  |  | Phenethyl benzoate | 7194 |  |
|  |  | Isocalamendiol | 12302240 |  |
|  |  | Butyl phthalate | 3026 |  |
|  |  | Hexadecanoic acid | 985 |  |
|  |  | Ferruginol | 442027 |  |
|  |  | Phytol | 5280435 |  |
|  |  | Hepta l | 8130 | [14] |
|  |  | Hexyl acetate | 8908 |  |
|  |  | Acetophenone | 7410 |  |
|  |  | Verbenone | 29025 |  |
|  |  | Pipero l | 8438 |  |
|  |  | Alpha-Copaene | 19725 |  |
|  |  | Cin maldehyde dimethyl acetal | 5463228 |  |
|  |  | Isoeugenol | 853433 |  |
|  |  | Thujopsene | 442402 |  |
|  |  | Coumarin | 323 |  |
| 7 | Gymnema sylvestre | 1,8-Cineole | 2758 | [15] |
|  |  | Octanol | 957 |  |
|  |  | Beta-Elemene | 6918391 |  |
|  |  | Acetophenone | 7410 |  |
|  |  | Germacrene A | 9548705 |  |
|  |  | p-Guaiacol | 9015 |  |
|  |  | Dodecanol | 8193 |  |
|  |  | Methyl eugenol | 7127 |  |
|  |  | 2-Pentadecanone | 61303 |  |
|  |  | 4-Ethyl guaiacol | 62465 |  |
|  |  | 2-Dodecenol | 90733 |  |
|  |  | Hexahydrofarnesyl acetone | 10408 |  |
|  |  | Eugenol | 3314 |  |
|  |  | m-Ethyl phenol | 12101 |  |
|  |  | Tetradecanol | 8209 |  |
|  |  | 4-Vinyl guaiacol | 332 |  |
|  |  | Methyl palmitate | 8181 |  |
|  |  | Ethyl palmitate | 12366 |  |
|  |  | Pentadecanol | 12397 |  |
|  |  | Tetradecadiene | 6365430 |  |
|  |  | 9,12,15-Octadecatrie l | 5283384 |  |
|  |  | 1-Hexadecanol | 2682 |  |
|  |  | Indole | 798 |  |
|  |  | Tetracosane | 12592 |  |
|  |  | Ethyl octadec-9-enoate | 8123 |  |
|  |  | Pentacosane | 12406 |  |
|  |  | Methyl linole te | 5319706 |  |
|  |  | Octadecanol | 8221 |  |
|  |  | Phytol | 5280435 |  |
|  |  | Myristic acid | 11005 |  |
|  |  | Heptacosane | 11636 |  |
|  |  | Pentadecanoic acid | 13849 |  |
|  |  | Palmitic acid | 985 |  |
|  |  | Hydroquinone | 785 |  |
|  |  | Propane, 1,1-diethoxy- | 20858 | [16] |
|  |  | Catechol | 289 |  |
|  |  | 3-Methoxyacetophenone | 11460 |  |
|  |  | 2,3,5,6-Tetrafluoroanisole | 75351 |  |
|  |  | 1,2,3,4-Cyclohexanetetrol | 278584 |  |
|  |  | Tetradecanoic acid | 11005 |  |
|  |  | Bicyclo[2.2.1]heptane, 1,3,3-trimethyl- | 138701 |  |
|  |  | 6-Octen-1-ol, 3,7-dimethyl-, formate | 7778 |  |
|  |  | n-Hexadecanoic acid | 985 |  |
|  |  | Isophytol | 10453 |  |
|  |  | Alpha-Santoline alcohol | 565379 |  |
|  |  | Hexadecanoic acid, 2-hydroxy-1-(hydroxymethyl)ethyl ester | 123409 |  |
|  |  | Phthalic acid, di(hept-3-yl) ester | 6518413 |  |
|  |  | Spiro[cyclopropane-1,2'-[6.7]diazabicyclo[3.2.2]non-6-ene] | 556389 |  |
|  |  | Squalene | 638072 |  |
|  |  | Gamma-Tocopherol | 92729 |  |
|  |  | 1-Docosene | 74138 |  |
|  |  | Vitamin E | 14985 |  |
|  |  | Cholane-5,20(22)-diene-3b-phenoxy | 91742606 |  |
|  |  | Beta-Amyrin | 73145 |  |
|  |  | 1,6,10,14,18,22-Tetracosahexaen-3-ol, 2,6,10,15,19,23-hexamethyl-, (all-E)- | 5366014 |  |
|  |  | Stigmasterol | 5280794 |  |
|  |  | Alpha-Tocopherol-Beta-D-mannoside | 597057 |  |
|  |  | A'-Neogammacer-22(29)-ene | 92155 |  |
|  |  | Phytol, acetate | 6428538 |  |
|  |  | Cedrene-V6 | 605018 |  |
| 8 | Linum usitatissimum L. | Methyl tetradecanoate | 31284 | [17] |
|  |  | Pentadecanoic acid, methyl ester | 23518 |  |
|  |  | 7-Hexadecenoic acid, methyl ester, (Z)- | 5364431 |  |
|  |  | 9-Hexadecenoic acid, methyl ester, (Z)- | 643801 |  |
|  |  | Hexadecanoic acid, methyl ester | 8181 |  |
|  |  | cis-10-Heptadecenoic acid, methyl ester | 16219491 |  |
|  |  | Heptadecanoic acid, methyl ester | 15609 |  |
|  |  | 9-Octadecenoic acid (Z)-, methyl ester | 5364509 |  |
|  |  | 9-Octadecenoic acid (Z)-, methyl ester | 5364509 |  |
|  |  | 9,12,15-Octadecatrienoic acid, methyl ester, (Z,Z,Z)- | 5319706 |  |
|  |  | Methyl stearate | 8201 |  |
|  |  | 11,14-Eicosadienoic acid, methyl ester | 5365566 |  |
|  |  | 11,14,17-Eicosatrienoic acid, methyl ester | 5367326 |  |
|  |  | cis-11-Eicosenoic acid, methyl ester | 5463047 |  |
|  |  | 8,11,14-Eicosatrienoic acid, methyl ester | 5366854 |  |
|  |  | Eicosanoic acid, methyl ester | 14259 |  |
|  |  | 13-Docosenoic acid, methyl ester, (Z)- | 5364423 |  |
|  |  | Docosanoic acid, methyl ester | 13584 |  |
|  |  | Octadecanoic acid, 6-hydroxy-, methyl ester | 559013 |  |
|  |  | 15-Tetracosenoic acid, methyl ester, (Z)- | 5364841 |  |
|  |  | Methyl 8-oxohexadecanoate | 546395 |  |
|  |  | Tetracosanoic acid, methyl ester | 75546 |  |
|  |  | Erucic acid | 5281116 |  |
|  |  | Hexacosanoic acid, methyl ester | 22048 |  |
|  |  | Stigmast-5-en-3-ol, oleate | 20831071 |  |
|  |  | 9,19-Cyclolanost-23-ene-3,25-diol, 3-acetate, (3.beta.,23E)- | 5363281 |  |
|  |  | Cholesterol | 5997 | [18] |
|  |  | Brassicasterol | 5281327 |  |
|  |  | Campesterol | 173183 |  |
|  |  | Stigmasterol | 5280794 |  |
|  |  | Beta-Sitosterol | 222284 |  |
|  |  | Obtusifoliol | 65252 |  |
|  |  | Cycloartenol | 92110 |  |
| 9 | Myrisrtica fragrance Houtt | Alpha-pinene | 6654 | [19] |
|  |  | Beta-phellandrene | 11142 |  |
|  |  | Terpinene | 92234 |  |
|  |  | Terpinen-4-ol | 11230 |  |
|  |  | Safrole | 5144 |  |
|  |  | Myristicin | 4276 |  |
|  |  | Alpha-Thujene | 17868 | [20] |
|  |  | Alpha-Pinene | 6654 |  |
|  |  | Camphene | 6616 |  |
|  |  | Beta-Pinene | 14896 |  |
|  |  | Sabinene | 18818 |  |
|  |  | Myrcene | 31253 |  |
|  |  | Delta-3-Carene | 26049 |  |
|  |  | Alpha-Phellandrene | 7460 |  |
|  |  | Alpha-Terpinene | 7462 |  |
|  |  | p-Cymene | 7463 |  |
|  |  | Limonene | 22311 |  |
|  |  | 1,8-Cineole | 2758 |  |
|  |  | Gamma-Terpinene | 7461 |  |
|  |  | Trans-Sabinene Hydrate | 6326181 |  |
|  |  | Terpinolene | 11463 |  |
|  |  | Li lool | 6549 |  |
|  |  | (Z)-p-Menth-2-en-1-ol | 13918681 |  |
|  |  | (E)-p-Menth-2-en-1-ol | 122484 |  |
|  |  | Terpinen-4-ol | 11230 |  |
|  |  | Alpha-Terpineol | 17100 |  |
|  |  | Carvacrol | 10364 |  |
|  |  | Safrole | 5144 |  |
|  |  | Eugenol | 3314 |  |
|  |  | Myristic Acid | 11005 |  |
|  |  | Geranyl Acetate | 1549026 |  |
|  |  | Isoeugenol | 853433 |  |
|  |  | Beta-Caryophyllene | 5281515 |  |
|  |  | Beta-Cubebene | 93081 |  |
|  |  | Trans-Alpha-Bergamotene | 6429302 |  |
|  |  | Germacrene D | 5317570 |  |
|  |  | Myristicin | 4276 |  |
|  |  | Elemicin | 10248 |  |
|  |  | Alpha-Thujene | 17868 | [21] |
|  |  | Alpha-Pinene | 6654 |  |
|  |  | Camphene | 6616 |  |
|  |  | 3-Carene | 26049 |  |
|  |  | Alpha-Phellandrene | 7460 |  |
|  |  | Beta-Myrcene | 31253 |  |
|  |  | 2-Carene | 79044 |  |
|  |  | Sabinene | 18818 |  |
|  |  | Beta-Pinene | 14896 |  |
|  |  | Myristicin | 4276 |  |
|  |  | Gamma-Terpinene | 7461 |  |
|  |  | Alpha-Terpinene | 7462 |  |
|  |  | 3,7,7-Trimethylcyclohepta-1,3,5-triene | 576718 |  |
|  |  | Limonene | 22311 |  |
|  |  | Isoterpinolene | 102443 |  |
|  |  | Gamma-Terpineol | 11467 |  |
|  |  | Alpha-Terpinolene | 11463 |  |
|  |  | Sylvestrene | 12304570 |  |
|  |  | Isomethyleugenol | 637776 |  |
|  |  | Cis-p-Menth-2-en-1-ol | 5319367 |  |
|  |  | 4-Propenyl Syringol | 176439 |  |
|  |  | Citronellyl Decanoate | 57353225 |  |
|  |  | Bicyclogermacrene | 13894537 |  |
|  |  | 4-Terpineol | 11230 |  |
|  |  | Cubebol | 11276107 |  |
|  |  | Cubene | 57178215 |  |
|  |  | Piperitol | 10247670 |  |
|  |  | Isoelemicin | 5318557 |  |
|  |  | Copaene | 12303902 |  |
|  |  | Beta-Copaene | 57339298 |  |
|  |  | Gamma-Amorphene | 12313019 |  |
|  |  | Cis-Alpha-Bergamotene | 6429303 |  |
|  |  | Isogermacrene | 91749707 |  |
|  |  | Licarin B | 6441061 |  |
|  |  | Bergamotene | 521569 |  |
|  |  | Gamma-Asarone | 636750 |  |
|  |  | Elemicin | 10248 |  |
|  |  | Myrislig n | 21636106 |  |
|  |  | Alpha-Thujene | 17868 | [22] |
|  |  | Alpha-Pinene | 6654 |  |
|  |  | Camphene | 6616 |  |
|  |  | Sabinene | 18818 |  |
|  |  | Beta-Pinene | 14896 |  |
|  |  | Beta-Myrcene | 31253 |  |
|  |  | Alpha-Terpinene | 7462 |  |
|  |  | Limonene | 22311 |  |
|  |  | Gamma-Terpinene | 7461 |  |
|  |  | Cis-Sabinenehydrate | 62367 |  |
|  |  | Alpha-Terpinolene | 11463 |  |
|  |  | Beta-Ocimene | 18756 |  |
|  |  | 3-Cyclohexen-1-ol | 556685 |  |
|  |  | Alpha-Cubebene | 442359 |  |
|  |  | Alpha-Copaene | 19725 |  |
|  |  | Trans-methyl isoeugenol | 1549045 |  |
|  |  | Trans-caryophyllene | 5281515 |  |
|  |  | Trans-Alpha-bergamotene | 6429302 |  |
|  |  | Alpha-Humulene | 5281520 |  |
|  |  | Germacrene | 9548705 |  |
|  |  | Gamma-Cadinene | 92313 |  |
|  |  | Myristicin | 4276 |  |
|  |  | Alpha-Thujene | 17868 | [23] |
|  |  | Alpha-Pinene | 6654 |  |
|  |  | Camphene | 6616 |  |
|  |  | Sabinene | 18818 |  |
|  |  | Beta-Pinene | 14896 |  |
|  |  | Isolimonene | 521268 |  |
|  |  | Beta-Myrcene | 31253 |  |
|  |  | Alpha-Phellandrene | 7460 |  |
|  |  | 3-Carene | 26049 |  |
|  |  | Alpha-Terpinene | 7462 |  |
|  |  | p-Cymene | 7463 |  |
|  |  | (S)-Limonene | 439250 |  |
|  |  | Trans-Beta-Ocimene | 5281553 |  |
|  |  | Gamma-Terpinene | 7461 |  |
|  |  | Cis-4-Thujanol | 12315152 |  |
|  |  | Terpinolene | 11463 |  |
|  |  | Li lool | 6549 |  |
|  |  | Alpha-Terpineol | 17100 |  |
|  |  | Cis-Pinocamphone | 84532 |  |
|  |  | Terpinen-4-ol | 11230 |  |
|  |  | Alpha,Alpha,4-Trimethylbenzenemethanol | 14529 |  |
|  |  | Alpha-Terpineol | 17100 |  |
|  |  | Cis-Piperitol | 85567 |  |
|  |  | Trans-Piperitol | 85568 |  |
|  |  | Cis-Sabinene Hydrate Acetate | 6427493 |  |
|  |  | Ascaridole | 10545 |  |
|  |  | Trans-Sabinene Hydrate Acetate | 6427504 |  |
|  |  | Safrole | 5144 |  |
|  |  | 2-Ethyl-4,5-Dimethylphenol | 247477 |  |
|  |  | Isoascaridole | 12314661 |  |
|  |  | Myrtenyl Acetate | 61262 |  |
|  |  | Delta-Elemene | 12309449 |  |
|  |  | Alpha-Terpinyl Acetate | 111037 |  |
|  |  | Alpha-Cubebene | 442359 |  |
|  |  | Eugenol | 3314 |  |
|  |  | Alpha-Copaene | 19725 |  |
|  |  | Methyl Eugenol | 7127 |  |
|  |  | Trans-Caryophyllene | 5281515 |  |
|  |  | Trans-Alpha-Bergamotene | 6429302 |  |
|  |  | Isoeugenol | 853433 |  |
|  |  | Beta-Ylangene | 25244198 |  |
|  |  | Germacrene D | 5317570 |  |
|  |  | Methyl Isoeugenol | 637776 |  |
| 10 | Nigella sativa L. | Beta-Pinene | 14896 | [24] |
|  |  | D-Glucose | 5793 |  |
|  |  | O-Cymene | 10703 |  |
|  |  | DL-Arabinose | 854 |  |
|  |  | Trans-4-methoxy thujane | 129845680 |  |
|  |  | 2-Propyl-tetrahydropyran-3-ol | 541755 |  |
|  |  | Terpinen-4-ol | 11230 |  |
|  |  | Alpha-D-Glucopyranoside | 9898327 |  |
|  |  | Thymoquinone | 10281 |  |
|  |  | 2-Isopropylidene-5-methylhex-4-e l | 534886 |  |
|  |  | Limonen-6-ol, pivalate | 545235 |  |
|  |  | Longifolene | 289151 |  |
|  |  | 2-(4-Nitrobutyryl)cyclooctanone | 544401 |  |
|  |  | Beta-Bisabolene | 10104370 |  |
|  |  | 1,1-Diphenyl-4-phenylthiobut-3-en-1-ol | 5363672 |  |
|  |  | l-(+)-Ascorbic acid 2,6-dihexadecanoate | 54722209 |  |
|  |  | 9,12-Octadecadienoic acid (Z,Z)-, methyl ester | 5284421 |  |
|  |  | 1-Heptatriacotanol | 537071 |  |
|  |  | 10,13-Eicosadienoic acid, methyl ester | 5365687 |  |
|  |  | E,E,Z-1,3,12-No decatriene-5,14-diol | 5364768 |  |
|  |  | Phthalic acid, decyl oct-3-yl ester | 19017886 |  |
|  |  | 1,2-Benzenedicarboxylic acid, bis(8-methylnonyl) ester | 33599 |  |
|  |  | Stigmasterol | 5280794 |  |
|  |  | 2-Chloroethyl vinyl sulfide | 157735 | [25] |
|  |  | 11-Octadecenoic acid, methyl ester | 5364432 |  |
|  |  | Cyclononene | 5463153 |  |
|  |  | Undec-10-ynoic acid, undecyl ester | 91692431 |  |
|  |  | Undec-10-ynoic acid, dodecyl ester | 91692432 |  |
|  |  | 9,12-Octadecadie l | 5283383 |  |
|  |  | n-Propyl 9,12-octadecadienoate | 9949088 |  |
|  |  | Propa mide | 6578 |  |
|  |  | Octan-2-one | 8093 |  |
|  |  | 9-Oxabicyclo[6.1.0]no ne | 67513 |  |
|  |  | Cyclopentaneundecanoic acid | 534549 |  |
|  |  | 4-Decyne | 16944 |  |
|  |  | 7-Hydroxy-3-(1,1-dimethylprop-2-enyl)coumarin | 5363192 |  |
|  |  | 2,6-Pyridinediamine | 8861 |  |
|  |  | 1-Nonylcycloheptane | 21262075 |  |
|  |  | 7,11-Hexadecadie l | 543335 |  |
|  |  | Oleic Acid | 445639 |  |
|  |  | cis-13-Octadecenoic acid | 5312441 |  |
|  |  | 9-Octadece l | 5283381 |  |
|  |  | Trimethylsilyl-di(timethylsiloxy)-silane | 6329086 |  |
|  |  | 3-Octyne, 6-methyl- | 534253 |  |
|  |  | Thymoquinone | 10281 | [26] |
|  |  | Dithymoquinone | 398941 |  |
|  |  | Alpha-hederin | 73296 |  |
|  |  | Carvacrol | 10364 |  |
|  |  | Thymol | 6989 |  |
|  |  | P-cymene | 7463 |  |
|  |  | Camphene | 6616 |  |
|  |  | Nigellidine | 136828302 |  |
|  |  | Nigellimine | 20725 |  |
|  |  | Nigellimine-N-oxide | 69131015 |  |
|  |  | Nigellidine | 136828302 |  |
|  |  | Linoleic Acid | 5280450 |  |
|  |  | Palmitic Acid | 985 |  |
|  |  | Beta-sitosterol | 222284 |  |
|  |  | Myristic Acid | 11005 |  |
|  |  | Arachidonic Acid | 444899 |  |
|  |  | Oleic Acid | 445639 |  |
|  |  | Gamma-linolenic Acid | 5280933 |  |
|  |  | Eugenol | 3314 |  |
|  |  | Alpha-Tocopherol | 14985 |  |
|  |  | Gamma-Tocopherol | 92729 |  |
|  |  | Beta-Pinene | 14896 | [27] |
|  |  | O-Cymene | 10703 |  |
|  |  | Thymoquinone | 10281 |  |
|  |  | Palmitic acid | 985 |  |
|  |  | Oleic acid | 445639 |  |
|  |  | Linoleic acid | 5280450 |  |
|  |  | 1-Heptatriacotanol | 537071 |  |
|  |  | Thymol | 6989 |  |
| 11 | Nyctanthes arbortristis L. | Octacosane | 12408 | [28] |
|  |  | Furfural | 7362 |  |
|  |  | Phytane | 12523 |  |
|  |  | Eicosamethyl-cyclodecasiloxane | 519601 |  |
|  |  | 1-Hexacosanol | 68171 |  |
|  |  | Oxirane | 6354 |  |
|  |  | 2-Methylbenzoic acid | 8373 |  |
|  |  | Furfural | 7362 | [29] |
|  |  | trans-2-Hexe l | 5281168 |  |
|  |  | 4-Cyclopentene-1,3-dione | 70258 |  |
|  |  | Benzaldehyde | 240 |  |
|  |  | Benzyl alcohol | 244 |  |
|  |  | Benzoic acid, methyl ester | 7150 |  |
|  |  | Phenylethyl alcohol | 6054 |  |
|  |  | Oxopholone | 62374 |  |
|  |  | Epoxyli lol | 26396 |  |
|  |  | Li lool oxide | 22310 |  |
|  |  | Safra l | 61041 |  |
|  |  | Eucarvone | 136330 |  |
|  |  | p-Vinylguaiacol | 332 |  |
|  |  | Methyl anthranilate | 8635 |  |
|  |  | Cin mic acid | 444539 |  |
|  |  | Hexahydrofarnesyl acetone | 10408 |  |
|  |  | Methyl hexadecanoate | 8181 |  |
|  |  | Methyl linoleate | 5284421 |  |
|  |  | Methyl elaidate | 5280590 |  |
|  |  | Phytol | 5280435 |  |
|  |  | Docosane | 12405 |  |
|  |  | Phytyl acetate | 637195 |  |
|  |  | Muscalure | 5365075 |  |
|  |  | Tricosane | 12534 |  |
|  |  | Pentacosane | 12406 |  |
|  |  | Heptacosane | 11636 |  |
|  |  | Supraene | 638072 |  |
|  |  | 2-Methyloctacosane | 519147 |  |
|  |  | No cosane | 12409 |  |
|  |  | Triacontane | 12535 |  |
| 12 | Ocimum tenuiflorum L. | Phenol, 2-methoxy-4-(2-propenyl)- | 3314 | [30] |
|  |  | Guanosine | 135398635 |  |
|  |  | Neophytadiene | 10446 |  |
|  |  | Hexadecanoic acid | 985 |  |
|  |  | Octanoic acid, 2-dimethylaminoethyl ester | 3075918 |  |
|  |  | 3-cyclopentylpropionic acid, 2-dimethylaminoethyl ester | 91693811 |  |
|  |  | Tofisopam | 5502 |  |
|  |  | Butanoic acid, ethyl ester | 7762 |  |
|  |  | Propane, 1-(1-ethoxyethoxy)- | 30220 |  |
|  |  | 1-butanol, 3-methyl-, acetate | 31276 |  |
|  |  | 4h-pyran-4-one, 2,3-dihydro-3,5-dihydroxy-6-methyl- | 119838 |  |
|  |  | Guanosine | 135398635 |  |
|  |  | 4-((1e)-3-hydroxy-1-propenyl)-2-methoxyphenol | 1549095 |  |
|  |  | Neophytadiene | 10446 |  |
|  |  | N-hexadecanoic acid | 985 |  |
|  |  | Phytol | 5280435 |  |
|  |  | Phytol, acetate | 6428538 |  |
|  |  | 3-cyclopentylpropionic acid, 2-dimethylaminoethyl ester | 91693811 |  |
|  |  | Propionic acid, 2-mercapto-, allyl ester | 548368 |  |
|  |  | Oxirane, 2,3-dimethyl-, cis- | 92162 |  |
|  |  | Propane, 1,1-diethoxy- | 20858 |  |
|  |  | Propane, 1-(1-ethoxyethoxy)- | 30220 |  |
|  |  | Mepronil | 41632 |  |
|  |  | 7-methoxy-3-(p-methoxyphenyl)-4h-chromen-4-one | 136419 |  |
|  |  | Cyclopropylmethanol | 75644 |  |
|  |  | 2,5-dihydro-1h-pyrrole | 66059 |  |
|  |  | Alpha-Pinene | 6654 | [31] |
|  |  | Camphene | 6616 |  |
|  |  | Sabinene | 18818 |  |
|  |  | Beta-Pinene | 14896 |  |
|  |  | Myrcene | 31253 |  |
|  |  | p-Cymene | 7463 |  |
|  |  | Limonene | 22311 |  |
|  |  | 1,8-Cineole | 2758 |  |
|  |  | (E)-Beta-Ocimene | 5281553 |  |
|  |  | (E)-Li lool oxide | 6432254 |  |
|  |  | Li lool | 6549 |  |
|  |  | Camphor | 2537 |  |
|  |  | Borneol | 64685 |  |
|  |  | Alpha-Terpineol | 17100 |  |
|  |  | Methyl chavicol | 8815 |  |
|  |  | Nerol | 643820 |  |
|  |  | Geraniol | 637566 |  |
|  |  | Eugenol | 3314 |  |
|  |  | (E)-Methyl cin mate | 637520 |  |
|  |  | Beta-Elemene | 6918391 |  |
|  |  | Methyl eugenol | 7127 |  |
|  |  | Beta-Caryophyllene | 5281515 |  |
|  |  | (E)-Alpha-Bergamotene | 6429302 |  |
|  |  | Alpha-Humulene | 5281520 |  |
|  |  | Germacrene D | 5317570 |  |
|  |  | Beta-Selinene | 442393 |  |
|  |  | Alpha-Selinene | 10856614 |  |
|  |  | Bicyclogermacrene | 13894537 |  |
|  |  | Beta-Bisabolene | 10104370 |  |
|  |  | Alpha-Bulnesene | 94275 |  |
|  |  | d-Cadinene | 441005 |  |
|  |  | Caryophyllene oxide | 1742210 |  |
| 13 | Punica gratum L. | Toluene | 1140 | [32] |
|  |  | Alpha-thujene | 17868 |  |
|  |  | Alpha-pinene, (-)- | 440968 |  |
|  |  | Camphene | 6616 |  |
|  |  | Sabinene | 18818 |  |
|  |  | Beta-myrcene | 31253 |  |
|  |  | 1-Phellandrene | 7460 |  |
|  |  | Delta-3-carene | 26049 |  |
|  |  | o-Cymene | 10703 |  |
|  |  | dl-Limonene | 22311 |  |
|  |  | Gamma-terpinene | 7461 |  |
|  |  | Alpha-terpinolene | 11463 |  |
|  |  | Alpha-terpinolene | 11463 |  |
|  |  | Camphor | 2537 |  |
|  |  | Cyclopentasiloxane, decamethyl- | 10913 |  |
|  |  | Alpha-amorphene | 12306046 |  |
|  |  | Trans-caryophyllene | 5281515 |  |
|  |  | 5-Acetamido-4,7-dioxo-4,7-dihydrobenzofurazan | 610143 |  |
|  |  | Dihydropyran | 8080 |  |
|  |  | Glycerin | 753 | [33] |
|  |  | Furfural | 7362 |  |
|  |  | Cyclobutylamine | 75645 |  |
|  |  | L-Glucose | 10954115 |  |
|  |  | Palmitic acid | 985 |  |
|  |  | 5-Hydroxymethylfurfural | 237332 |  |
|  |  | Heptasiloxane, hexadecamethyl- | 10912 |  |
|  |  | Octadecanoic acid | 5281 |  |
|  |  | Lanosterol | 246983 |  |
|  |  | Cycloartenol acetate | 17750996 |  |
|  |  | Ellagic acid | 5281855 | [34] |
|  |  | Gallic acid | 370 |  |
|  |  | Catechin | 9064 |  |
|  |  | Quercetin | 5280343 |  |
|  |  | Rutin | 5280805 |  |
|  |  | Cin mic acid | 444539 |  |
|  |  | Genistein | 5280961 |  |
|  |  | Kaempferol | 5280863 |  |
|  |  | Cyanidin | 128861 |  |
|  |  | Punicalin | 92131301 |  |
|  |  | Delphinidin | 68245 |  |
|  |  | Punicalagin | 16129719 |  |
|  |  | Linoleic acid | 5280450 |  |
|  |  | Chlorogenic acid | 1794427 |  |
|  |  | Luteolin | 5280445 |  |
| 14 | Saussurea costus (Falc.) Lipsch | Undecane | 14257 | [35] |
|  |  | Cyclohexane, (1-methylpropyl)- | 23468 |  |
|  |  | 2,3-Dimethyldecane | 86544 |  |
|  |  | Hexadecane | 11006 |  |
|  |  | 3-methyl-Undecane | 521960 |  |
|  |  | 4-methyl-1-Undecene | 522551 |  |
|  |  | Eicosene | 18936 |  |
|  |  | 5-methyl-Tetradecane | 98976 |  |
|  |  | 4-Methyldodecane | 521958 |  |
|  |  | 2,3-Dimethyldodecane | 521959 |  |
|  |  | Dodecane, 2,6,10-trimethyl- | 19773 | [36] |
|  |  | Hexadecane, 2,6,10,14-tetramethyl- | 12523 |  |
|  |  | Tetradecane | 12389 |  |
|  |  | 1,8,11-Heptadecatriene | 13573140 |  |
|  |  | 1,8,11,14-Heptadecatetraene | 5319559 |  |
|  |  | 1-Heptadecene | 23217 |  |
|  |  | 1-Octadecene | 8217 |  |
|  |  | 1-Docosene | 74138 |  |
|  |  | Octadecane | 11635 |  |
|  |  | No decane | 12401 |  |
|  |  | 1-Tricosene | 181154 |  |
|  |  | Eicosane | 8222 |  |
|  |  | 1-Tetracosene | 82436 |  |
|  |  | Heneicosane | 12403 |  |
|  |  | 1-No decene | 29075 |  |
|  |  | Styrene | 7501 |  |
|  |  | Alpha-Pinene | 6654 |  |
|  |  | Beta-Phellandrene | 11142 |  |
|  |  | Beta-Cymene | 10812 |  |
|  |  | Camphene | 6616 |  |
|  |  | Gamma-Terpinene | 7461 |  |
|  |  | Alpha-Phellandrene | 7460 |  |
|  |  | Beta-Elemene | 6918391 |  |
|  |  | Beta-Farnesene | 5281517 |  |
|  |  | Beta-Bisabolene | 10104370 |  |
|  |  | Caryophyllene | 5281515 |  |
|  |  | Alpha-Ionone | 5282108 |  |
|  |  | Alpha-Bergamotene | 86608 |  |
|  |  | Gamma-Ionone | 5363741 |  |
|  |  | Eudesma-4(14),11-diene | 442393 |  |
|  |  | Alpha-Guaiene | 5317844 |  |
|  |  | Alpha-Selinene | 10856614 |  |
|  |  | Beta-Humulene | 5318102 |  |
|  |  | Gamma-Bisabolene | 3033866 |  |
|  |  | Beta-Sesquiphellandrene | 12315492 |  |
|  |  | Alpha-Bisabolene | 86597 |  |
|  |  | Beta-Guaiene | 6949 |  |
|  |  | Alpha-Phellandrene, dimer | 6431154 |  |
|  |  | Alpha-Curcumene | 92139 |  |
|  |  | Gamma-Curcumene | 12304273 |  |
|  |  | Beta-Curcumene | 6428461 |  |
|  |  | Alpha-Bergamotol | 91749656 |  |
|  |  | Beta-Costol | 12304104 |  |
|  |  | Saussurea lactone | 556963 |  |
|  |  | Vanillosmin | 100572 |  |
|  |  | Dehydrocostus lactone | 73174 |  |
|  |  | Santamarine | 188297 |  |
|  |  | Reynosin | 482788 |  |
|  |  | Eucalyptol | 2758 |  |
|  |  | 13-Octadece l | 5367670 |  |
|  |  | 3,7,11-trimethyl-1-Dodecanol | 138824 |  |
|  |  | 2-Isopropyl-5-methyl-1-heptanol | 545941 |  |
|  |  | Pentadecanol | 12397 |  |
|  |  | Nerolidol | 5284507 |  |
|  |  | Alpha-Eudesmol | 92762 |  |
|  |  | Beta-Eudesmol | 91457 |  |
|  |  | Vulgarol B | 91748781 |  |
|  |  | Valerenol | 91699505 |  |
|  |  | 1-Heneicosanol | 85014 |  |
|  |  | 1-Heptacosanol | 74822 |  |
|  |  | Octacosanol | 68406 |  |
|  |  | Humulenol-II | 102115341 |  |
|  |  | Valerenyl acetate | 91747208 |  |
|  |  | Palmitic acid, methyl ester | 8181 |  |
|  |  | Methyl stearate | 8201 |  |
|  |  | Tributyl acetylcitrate | 6505 |  |
|  |  | Lupeol | 259846 |  |
|  |  | Beta-Sitosterol acetate | 5354503 |  |
|  |  | Beta-Sitosterol | 222284 |  |
|  |  | Docosahexaenoic acid | 445580 |  |
|  |  | Caproaldehyde | 6184 | [37] |
|  |  | Furfural | 7362 |  |
|  |  | Alpha-Thujene | 17868 |  |
|  |  | Alpha-Pinene | 6654 |  |
|  |  | Camphene | 6616 |  |
|  |  | Beta-Pinene | 14896 |  |
|  |  | Myrcene | 31253 |  |
|  |  | Sabinene | 18818 |  |
|  |  | Alpha-Terpinene | 7462 |  |
|  |  | p-Cymene | 7463 |  |
|  |  | Limonene | 22311 |  |
|  |  | Beta-Phellandrene | 11142 |  |
|  |  | 1,8 Cineol | 2758 |  |
|  |  | (E)-Beta-Ocimene | 5281553 |  |
|  |  | Gamma-Terpinene | 7461 |  |
|  |  | Alpha-Terpinolene | 11463 |  |
|  |  | Li lool | 6549 |  |
|  |  | Chrysanthenone | 442463 |  |
|  |  | Camphor | 2537 |  |
|  |  | Menthone | 26447 |  |
|  |  | Alpha-Fenchene | 28930 |  |
|  |  | Citronellal | 7794 |  |
|  |  | Terpinen-4-ol | 11230 |  |
|  |  | Cryptone | 92780 |  |
|  |  | Alpha-Terpineol | 17100 |  |
|  |  | Estragole | 8815 |  |
|  |  | Preg ne | 6857422 |  |
|  |  | Anethol | 637563 |  |
|  |  | Thymol | 6989 |  |
|  |  | Citronellyl Propio te | 8834 |  |
|  |  | Alpha-Copaene | 19725 |  |
|  |  | Beta-Elemene | 6918391 |  |
|  |  | Alpha-Cederene | 442348 |  |
|  |  | Alpha-Ionone | 5282108 |  |
|  |  | (E)-Caryophyllene | 5281515 |  |
|  |  | Alpha-Humulene | 5281520 |  |
|  |  | Geranyl Acetone | 1549778 |  |
|  |  | Allo-Aromadendrene | 42608158 |  |
|  |  | Beta-Selinene | 442393 |  |
|  |  | Alpha-Curcumene | 92139 |  |
|  |  | Beta-Ionone | 638014 |  |
|  |  | Alpha-Selinene | 10856614 |  |
|  |  | Beta-Guaiene | 6949 |  |
|  |  | Beta-Himachalene | 11586487 |  |
|  |  | Delta-Cadinene | 441005 |  |
|  |  | Cis-Alpha-Bisabolene | 5352653 |  |
|  |  | Elemol | 92138 |  |
|  |  | Nerolidol | 5284507 |  |
|  |  | Caryophyllene Oxide | 1742210 |  |
|  |  | (E,Z)-Alpha-Farnesene | 442368 |  |
|  |  | Beta-Eudesmol | 91457 |  |
|  |  | Alpha-Eudesmol | 92762 |  |
|  |  | Heneicosane | 12403 |  |
|  |  | Vulgarol B | 91748781 |  |
|  |  | Valerenol | 91699505 |  |
|  |  | Dehydrocostus Lactone | 73174 |  |
|  |  | Dehydrosaussurea Lactone | 556920 |  |
|  |  | Methyl Linoleate | 5284421 |  |
|  |  | Ethyl Linoleate | 5282184 |  |
|  |  | Anethole | 637563 | [38] |
|  |  | Thymol | 6989 |  |
|  |  | Citronellyl propio te | 8834 |  |
|  |  | Alpha-Thujene | 17868 |  |
|  |  | Alpha-Pinene | 6654 |  |
|  |  | Camphene | 6616 |  |
|  |  | Beta-Pinene | 14896 |  |
|  |  | Camphor | 2537 |  |
|  |  | Myrcene | 31253 |  |
|  |  | Sabinene | 18818 |  |
|  |  | p-Cymene | 7463 |  |
|  |  | Limonene | 22311 |  |
|  |  | Gamma-Terpinene | 7461 |  |
|  |  | Li lool | 6549 |  |
|  |  | Menthone | 26447 |  |
|  |  | Citronellal | 7794 |  |
|  |  | Terpinen-4-ol | 11230 |  |
|  |  | Alpha-Terpineol | 17100 |  |
|  |  | Estragole | 8815 |  |
|  |  | Alpha-Terpinolene | 11463 |  |
|  |  | Dehydrocostus lactone | 73174 |  |
|  |  | Zaluzanin C | 72646 |  |
|  |  | Cy ropicrin | 119093 |  |
|  |  | Lappalone | 10979139 |  |
|  |  | Mokko lactone | 167495 |  |
|  |  | Hexadecane | 11006 |  |
|  |  | Octacosane, 1-Iodo | 12696144 |  |
|  |  | Hentriacontane | 12410 |  |
|  |  | Heptadecane, 2,6,10,15-Tetramethyl | 41209 |  |
|  |  | Tritetracontane | 522398 |  |
|  |  | Carbonic acid, decyl undecyl ester | 91693139 |  |
|  |  | 6-Tetradecanesulfonic acid, butyl ester | 551402 |  |
|  |  | Heneicosane | 12403 |  |
|  |  | Dotriacontane | 11008 |  |
|  |  | Seli -3,7(11)-diene | 6432648 |  |
| 15 | Syzygium aromaticum (L.) | 2-Nonanone | 13187 | [39] |
|  |  | (E)-4,8-Dimethyl-1,3,7-no triene | 6427110 |  |
|  |  | Acetic acid, phenylmethyl ester | 8785 |  |
|  |  | Methyl salicylate | 4133 |  |
|  |  | Chavicol | 68148 |  |
|  |  | Alpha-Cubebene | 442359 |  |
|  |  | Eugenol | 3314 |  |
|  |  | Alpha-Copaene | 19725 |  |
|  |  | Cis-isoeugenol | 1549041 |  |
|  |  | Beta-Elemene | 6918391 |  |
|  |  | Caryophyllene | 5281515 |  |
|  |  | Beta-Gurjunene | 6450812 |  |
|  |  | Alpha-Ylangene | 442409 |  |
|  |  | Alpha-Humulene | 5281520 |  |
|  |  | Alloaromadendrene | 10899740 |  |
|  |  | Delta-Cadinene | 441005 |  |
|  |  | Gamma-Muurolene | 12313020 |  |
|  |  | Alpha-Amorphene | 12306046 |  |
|  |  | Alpha-Muurolene | 12306047 |  |
|  |  | Beta-Selinene | 442393 |  |
|  |  | Alpha-Selinene | 10856614 |  |
|  |  | Beta-Cadinene | 10657 |  |
|  |  | Gamma-Cadinene | 92313 |  |
|  |  | Eugenyl acetate | 7136 |  |
|  |  | Cadi -1,4-diene | 6427091 |  |
|  |  | Alpha-Cadinene | 12306048 |  |
|  |  | Alpha-Calacorene | 12302243 |  |
|  |  | 1,Z-5,E-7-Dodecatriene | 5367454 |  |
|  |  | Caryophyllenyl alcohol | 91704770 |  |
|  |  | (-)-Caryophyllene oxide | 1742210 |  |
|  |  | Junipene | 1796220 |  |
|  |  | Ledol | 92812 |  |
|  |  | Humulene Oxide | 6324 |  |
|  |  | Beta-Guaiene | 6949 |  |
|  |  | Tau-Muurolol | 6432221 |  |
|  |  | Vulgarol B | 91748781 |  |
|  |  | 2,3,4-Trimethoxyacetophenone | 83810 |  |
|  |  | Benzyl benzoate | 2345 |  |
|  |  | 2-Heptanone | 8051 | [40] |
|  |  | Alpha-Pinene | 6654 |  |
|  |  | Sabinene | 18818 |  |
|  |  | Myrcene | 31253 |  |
|  |  | Alpha-Phellandrene | 7460 |  |
|  |  | Ethyl-Hexanoate | 31265 |  |
|  |  | Alpha-Terpinene | 7462 |  |
|  |  | 1,8-Cineole | 2758 |  |
|  |  | D-Limonene | 440917 |  |
|  |  | p-Cymene | 7463 |  |
|  |  | Gamma-Terpinene | 7461 |  |
|  |  | 2-No none | 13187 |  |
|  |  | Terpinolene | 11463 |  |
|  |  | Li lool | 6549 |  |
|  |  | Benzyl acetate | 8785 |  |
|  |  | Benzoic acid, ethyl ester | 7165 |  |
|  |  | Terpinen-4-ol | 11230 |  |
|  |  | Alpha-Terpineol | 17100 |  |
|  |  | Chavicol | 68148 |  |
|  |  | Geraniol | 637566 |  |
|  |  | Alpha-Terpinyl acetate | 111037 |  |
|  |  | Eugenol | 3314 |  |
|  |  | Geranyl acetate | 1549026 |  |
|  |  | Beta-Caryophyllene | 5281515 |  |
|  |  | Humulene | 5281520 |  |
|  |  | Gamma-Cadinene | 92313 |  |
|  |  | Eugenol acetate | 7136 |  |
|  |  | Germacrene B | 5281519 |  |
|  |  | Caryophyllene oxide | 1742210 |  |
|  |  | Alpha-Amorphene | 12306046 |  |
|  |  | Gamma-Muurolene | 12313020 |  |
|  |  | Beta-Selinene | 442393 |  |
|  |  | Alpha-Selinene | 10856614 |  |
|  |  | Alpha-Farnesene | 5281516 |  |
|  |  | Methyl salicylate | 4133 | [41] |
|  |  | Chavicol | 68148 |  |
|  |  | (E)-Cin maldehyde | 637511 |  |
|  |  | Eugenol | 3314 |  |
|  |  | Alpha-Humulene | 5281520 |  |
|  |  | 9-epi-(E)-Caryophyllene | 6429274 |  |
|  |  | Eugenol acetate | 7136 |  |
|  |  | (Z)-Lanceol | 15560069 |  |
| 16 | Trigonella foenum-graecum L. | 2-Methylpyrrolidine | 13003 | [42] |
|  |  | Dodecane | 8182 |  |
|  |  | Alpha-Terpinene | 7462 |  |
|  |  | Alpha-Terpinyl acetate | 111037 |  |
|  |  | Tetradecane | 12389 |  |
|  |  | Caryophyllene | 5281515 |  |
|  |  | Pentadecane | 12391 |  |
|  |  | Trichloroacetic acid, pentadecyl ester | 522535 |  |
|  |  | Pinene | 15837102 |  |
|  |  | Phytol | 5280435 |  |
|  |  | No decane | 12401 |  |
|  |  | Eicosane | 8222 |  |
|  |  | Linoleic acid methyl ester | 5284421 |  |
|  |  | Linoleic acid | 5280450 |  |
|  |  | 4-Pentyl-1-(4-propylcyclohexyl)-1-cyclohexene | 557007 |  |
|  |  | 1-Piperidinepropanenitrile | 18338 |  |
|  |  | Palmidrol | 4671 |  |
|  |  | Glyceryl 2-linoleate | 5365676 |  |
|  |  | (R)-(-)-(Z)-14-Methyl-8-hexadecen-1-ol | 12487634 |  |
|  |  | Quinoline | 7047 |  |
|  |  | Tetratriacontane | 26519 |  |
|  |  | Catechin 3-O-gallate | 5276454 | [43] |
|  |  | Sativanone | 13886678 |  |
|  |  | Gallic acid | 370 |  |
|  |  | Kampferol | 5280863 |  |
|  |  | 5-O-Feruloylquinic acid | 10133609 |  |
|  |  | Cyanidin | 128861 |  |
|  |  | Coumaroylquinic acid | 6441280 |  |
|  |  | Quercetin | 5280343 |  |
|  |  | Schisandrin C | 119112 |  |
|  |  | Pinoresinol | 73399 |  |
|  |  | Glycitin | 187808 |  |
|  |  | 3,5-Octadiene | 5352266 | [44] |
|  |  | p-Xylene | 7809 |  |
|  |  | Delta-3-Carene | 26049 |  |
|  |  | Hepta l | 8130 |  |
|  |  | 3-Isopropyltoluene | 10812 |  |
|  |  | Limonene | 22311 |  |
|  |  | Deca l | 8175 |  |
|  |  | 1-Methoxy-4-(2-propenyl)-benzene | 8815 |  |
|  |  | 2-Methyl-5-isopropylphenol | 10364 |  |
|  |  | Trans-anethole | 637563 |  |
|  |  | 5-Pentyl-2(5H)-furanone | 89559 |  |
|  |  | Cis-calamenene | 6429077 |  |
|  |  | Cadi -1,4-diene | 6427091 |  |
|  |  | Hexadecanoic acid | 985 |  |
|  |  | Beta-Thujone | 91456 |  |
| 17 | Ziziphus mauritia Lam. | 4-O-Caffeoylquinic acid | 9798666 | [45] |
|  |  | Caffeic acid | 689043 |  |
|  |  | Syringic acid | 10742 |  |
|  |  | Epicatechin | 72276 |  |
|  |  | p-Coumaric acid | 637542 |  |
|  |  | Quercitrin | 5280459 |  |
|  |  | Quercetin | 5280343 |  |
|  |  | ringenin | 439246 |  |
|  |  | Gallic acid | 370 |  |
|  |  | Squalene | 638072 |  |
|  |  | n-Hexadecanoic acid | 985 |  |
|  |  | Luteolin | 5280445 |  |
|  |  | Cirsilineol | 162464 |  |
|  |  | Tetradecanoic acid | 11005 |  |
|  |  | Hyperoside | 5281643 |  |
|  |  | Phthalic acid, butyl undecyl ester | 6423450 |  |
|  |  | Z-2-Dodecenol | 5364955 |  |
|  |  | L-(+)-Ascorbic acid 2,6-dihexadecanoate | 54722209 |  |
|  |  | Myristic acid | 11005 |  |
|  |  | Ferulic acid | 445858 |  |
|  |  | Betulinic acid | 64971 |  |
|  |  | D-Allose | 439507 |  |
|  |  | Linoleic acid | 5280450 |  |
|  |  | Alpha-Linolenic acid | 5280934 |  |
|  |  | Methyl palmitate | 8181 |  |
|  |  | Palmitic acid | 985 |  |
|  |  | Rutin | 5280805 |  |
|  |  | Methyl stearate | 8201 |  |
|  |  | Stearic acid | 5281 |  |
|  |  | Bacchotricuneatin C | 551422 |  |
|  |  | Lauric acid | 3893 |  |
|  |  | Vitamin E | 14985 |  |

[23]	Z. X. Zhao XiangSheng, W. H. Wu HaiFeng, W. J. Wei JianHe, and Y. M. Yang MeiHua, “Quantification and characterization of volatile constituents in Myristica fragrans Houtt. by gas chromatography-mass spectrometry and gas chromatography quadrupole-time-of-flight mass spectrometry.,” 2019.

[24]	M. Y. Hadi, G. J. Mohammed, and I. H. Hameed, “Analysis of bioactive chemical compounds of Nigella sativa using gas chromatography-mass spectrometry,” *Journal of Pharmacognosy and Phytotherapy*, vol. 8, no. 2, pp. 8–24, 2016.

[25]	B. C. Adebayo-Tayo, A. I. Briggs-Kamara, and A. M. Salaam, “Phytochemical composition, antioxidant, antimicrobial potential and gc-ms analysis of crude and partitioned fractions of Nigella sativa seed extract,” *Acta Microbiol. Bulg*, vol. 37, pp. 34–45, 2021.

[26]	M. Abbas *et al.*, “Antimicrobial Properties and Therapeutic Potential of Bioactive Compounds in Nigella sativa: A Review,” *Molecules*, vol. 29, no. 20, p. 4914, 2024.

[27]	F. A. Saleh, N. El-Darra, K. Raafat, and I. El Ghazzawi, “Phytochemical analysis of Nigella sativa L. Utilizing GC-MS exploring its antimicrobial effects against multidrug-resistant bacteria,” *Pharmacognosy Journal*, vol. 10, no. 1, 2018.

[28]	M. K. Shirsat, I. J. Singhvi, K. Gupta, and A. Garg, “IDENTIFICATION OF BIOACTIVE COMPOUNDS OF NYCTANTHES ARBORTRISTIS LINN BY GC-MS”.

[36]	A. Khan, A. H. Shah, and N. Ali, “In-vitro propagation and phytochemical profiling of a highly medicinal and endemic plant species of the himalayan region (Saussurea costus). Sci. Rep. 11, 23575,” 2021.

[37]	R. K. Nadda, A. Ali, R. C. Goyal, P. K. Khosla, and R. Goyal, “Aucklandia costus (syn. Saussurea costus): Ethnopharmacology of an endangered medicinal plant of the Himalayan region,” *J Ethnopharmacol*, vol. 263, p. 113199, 2020.

[38]	R. Kumari *et al.*, “Saussurea costus (Falc.) Lipsch.: a comprehensive review of its pharmacology, phytochemicals, ethnobotanical uses, and therapeutic potential,” *Naunyn Schmiedebergs Arch Pharmacol*, vol. 397, no. 3, pp. 1505–1524, 2024.

[39]	B. Amelia, E. Saepudin, A. H. Cahyana, D. U. Rahayu, A. S. Sulistyoningrum, and J. Haib, “GC-MS analysis of clove (Syzygium aromaticum) bud essential oil from Java and Manado,” in *AIP conference Proceedings*, AIP Publishing, 2017.
